# The mood stabilizer lithium alters behaviour and physiology via the gut brain axis

**DOI:** 10.64898/2026.01.26.701868

**Authors:** Daniel W. Thorpe, Yoichiro Ootsuka, Rochelle A. Peterson, Lauren A. Jones, Adam Humenick, Alyce M. Martin, Jett Zivkovic, Junichi Sakaguchi, Jack McArdle, Arne Ittner, Simon J. Brookes, Ali Habib, Rata Sirimaharaj, Youssef Tawodros, Michael Roach, Robert Edwards, Hans Clevers, Joep Beumer, Jens Puschof, Lai Wei, Rajan Singh, Se Eun Ha, Seungil Ro, Jasenka Zubcevic, Emily B. Otmanowski, Rebeca Mendez-Hernandez, Guillaume de Lartigue, William W. Blessing, Damien J. Keating

## Abstract

Lithium, introduced 75 years ago by John Cade^1^, remains the most effective mood stabilizer for bipolar disorder^2^. Lithium is proposed to modulate an array of cellular pathways, many ubiquitous to all cells, with pleiotropic roles unlinked to bipolar disorder or lithium responsiveness in genome-wide association studies^3,4^. These mechanisms cannot explain lithium’s specific effects on mood and behaviour. We demonstrate that lithium’s primary action is in the periphery, not in the brain itself. Lithium acts in the gut to trigger behavioural and physiological changes, akin to those associated with a torpor-like state, that protect individuals from ingested toxins. Lithium activates gastrointestinal enterochromaffin (EC) cells via their Trpm2 cation channels to modulate afferent vagal and area postrema inputs to the brain. Eliminating these inputs by focal brain lesions eliminates lithium’s effects, as does ablation of EC cells or their Trpm2 expression. Lithium’s Trpm2-dependent activation of EC cells also occurs in human gut tissue, providing translational relevance for our discovery. These findings challenge the prevailing perception that lithium acts directly on the brain. Via a previously unsuspected gut-brain pathway, lithium engages brain circuitry that reduces arousal and interaction with the external world, therapeutic goals in the manic phase of bipolar disorder.

---

The remarkable calming effect of lithium, the third element of the periodic table, was famously demonstrated by John Cade^1^, who observed that lithium-treated guinea pigs, “*although fully conscious, became extremely lethargic and unresponsive to stimuli for one to two hours before once again becoming normally active and timid*”. Today, 75 years later, lithium remains a first-line treatment for bipolar disorder, especially for stabilizing the manic phase and reducing suicide risk^2^. However, despite lithium’s clinical success, the mechanisms of its therapeutic action remain undiscovered. Proposed mechanisms have included inhibition of glycogen synthesis kinase-3β, modulation of oxidative metabolism, autophagy, neurogenesis and apoptosis^5^. However, these are ubiquitous cellular processes, not specific to any one brain region or neuronal subtype. They offer little insight into where lithium acts or how it selectively alters mood, arousal, and behaviour. This disconnect has been described as a paradox: *“a drug with such complex effects comes closest to having the most specific clinical effects in all of psychiatry”*^3^.

Evidence from animal studies already suggests that lithium’s calming effect may be initiated outside the brain. Systemic administration of lithium robustly reduces exploratory behaviours and aggression, induces conditioned taste aversions and substantially lowers body temperature^6–15^. These effects are not reproduced when lithium is delivered directly into the cerebrospinal fluid, suggesting that direct effects on the brain are insufficient. Moreover, these effects are abolished when communication between the gut and brain is disrupted by eliminating vagal afferent input to the brainstem via the nucleus tractus solitarius (NTS), and by lesioning the area postrema (AP), a chemosensory circumventricular organ that detects blood-borne signals in the brainstem^10,11,13,15,16^. Together, these findings raise the possibility that lithium’s therapeutic effects may be initiated via peripheral sensory pathways, rather than via direct central nervous system targets.

Lithium-induced nausea and vomiting has been regarded as “non-specific toxicity”^17^, yet these symptoms closely resemble coordinated defensive responses that protect the body from ingested toxins. Such gut-initiated brain-integrated responses can prevent absorption of toxin (nausea and vomiting), minimize physiological damage (via hypothermia and reduced metabolic rate), and deter future ingestion (through conditioned taste aversions and reduced exploratory activity). Notably, the early phase of nausea in humans is accompanied by behavioural withdrawal, reduced arousal and diminished impulsivity, changes aligned with therapeutic goals in the management of mania.

We therefore propose the novel hypothesis that lithium’s therapeutic action begins in the gut, activating specific brain pathways via the gut-brain axis. We focus on enterochromaffin (EC) cells, which synthesize and secrete serotonin (5-hydroxytryptamine, 5-HT) and particular neuropeptides^18–23^. Activation of EC cells could stimulate vagal afferent neurons and modulate the AP, providing two key routes for gut-derived signals to influence brain circuits mediating lithium’s behavioural and physiological effects. To test our hypothesis, we conducted experiments in guinea pigs, rats and mice, as well as ex vivo studies of human gut tissue, using amounts of lithium to emulate the therapeutic plasma range for bipolar disorder (0.5-1.2 mEq/L).

## Revisiting John Cade’s guinea pig lithium experiment

We began by reproducing Cade’s original observation that lithium induces marked lethargy in guinea pigs. At Cade’s reported dose of 6 mEq/kg, animals did not resist handling and showed a dose-dependent reduction in locomotor activity (Fig. 1a and Movie 1). Remarkably, we also found that lithium produced a dose-dependent drop in core body temperature (Fig 1b), almost 3°C after 6 mEq/kg. Within minutes of administration, most animals adopted a prone posture with at least one leg and paw splayed out. Paw temperature increased (Fig. 1c), consistent with enhanced peripheral blood flow. The extent of LiCl-induced hypothermia correlated with the time spent in the splayed posture (Fig. 1d). Similar reductions in body temperature were observed in rats, consistent with earlier reports^8^, and infrared imaging confirmed rapid dilatation of the thermoregulatory vessels in the tail within minutes of even low doses of lithium (Fig 1e).

**Fig. 1.**
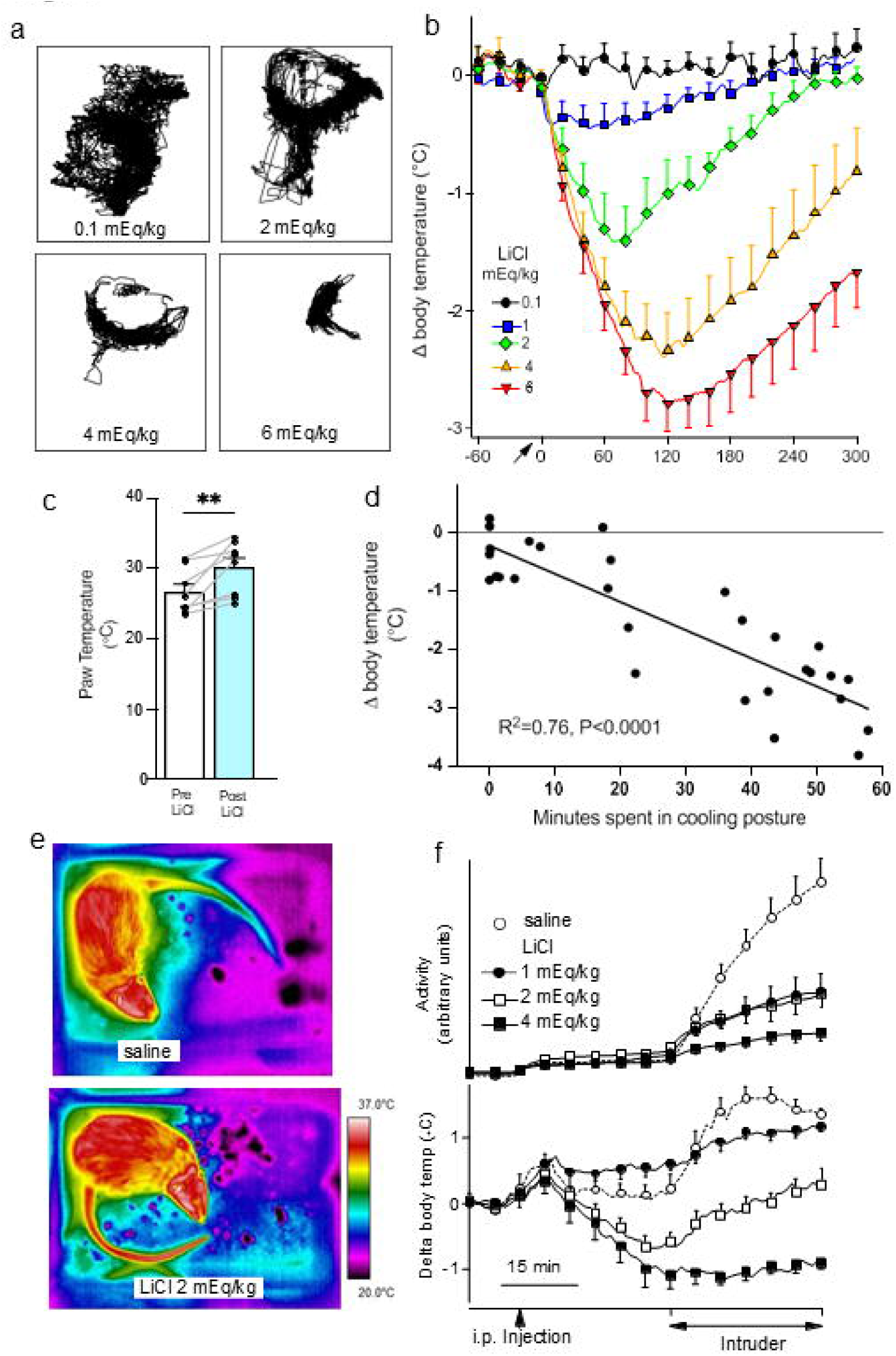
Lithium decreases activity and body temperature. **a** Video-based tracking of movement of an individual guinea pig 80-130 min after increasing doses of i.p. LiCl. **b** Body temperature in guinea pigs before and after lithium treatment (log-dose linear regression of minimum temperatures, R^2^=0.70, F=277.87, P<0.0001, n=6 animals per dose). **c** Guinea pig paw temperature before and 20 min after 4 mEq/kg i.p. LiCl **P<0.01, t =3.78, n=8). **d** Relation between LiCl-induced fall in body temperature and time spent prone in guinea pigs (linear regression R^2^=0.76, F=88.52, P<00001). **e** Infrared images of a rat showing increased tail temperature measured 6 minutes after LiCl or saline. **f** Dose-dependent (1, 2, 4 mEq/kg, n=6) LiCl inhibition of aggressive behavioural activity and rise in body temperature in resident rat elicited by introduction of an intruder rat (intruder-period log dose linear regression analysis, R^2^=0.85, P < 0.0001 and R^2^=0.52, P < 0.0001, respectively).

The results reveal that Cade’s “lethargy” reflects a coordinated behavioural and thermoregulatory response involving both reduced arousal and active heat dissipation. The splayed-out posture is best understood as “splooting” to dissipate heat rather than “lying on belly”, taken as an indication of illness^24^. In sickness, peripheral vasoconstriction limits heat loss^25^, whereas lithium causes peripheral vasodilation. Moreover, rats adopt the splayed-out posture in the absence of lithium in a conditioned taste aversion paradigm^24^, suggesting that it is part of a learned thermoregulatory response to a perceived toxin. It is notable that Cade’s guinea pigs all recovered to normality within several hours of lithium administration^1^.

By revisiting Cade’s work, we confirmed his “lethargy” observation and identified lithium-induced hypothermia and active heat dissipation as rapid, reproducible responses across species. Our findings place lithium’s behavioural effects in the broader context of a toxin-defense program, initiated in the gut and coordinated by the brain. Lithium-induced hypothermia may have protected Cade’s guinea pigs from the lethal consequence of systemically-administered urea solutions^1^, explaining his previously puzzling observation.

## Lithium reduces the aggressive response to an intruder animal in conscious rats

Lithium’s ability to reduce impulsivity, irritability, and aggression is a well-recognized clinical benefit in the treatment of mania^26,27^. To determine whether this effect also occurs in rodents, we used the resident-intruder paradigm, a standard model of aggression. Lithium dose-dependently reduced intruder provoked exploratory/aggressive behaviour in a resident rat, with substantial suppression occurring with a dose of 1 mEq/kg (Fig. 1f upper panel), a dose within the human therapeutic range. Lithium also dose-dependently attenuated emotional hyperthermia, the rise in body temperature that normally accompanies aggressive encounters^28^ (Fig. 1f lower panel). The coupling of reduced aggression with blunted emotional hyperthermia suggests a coordinated modulation of behavioural and autonomic outputs, consistent with engagement of specific brainstem circuits integrating gut sensory input. This dual effect supports our hypothesis that lithium acts via the gut-brain axis to influence both emotional state and physiological arousal.

## Lithium activates a gut-brain circuit

To identify lithium-activated gut-brain circuits in conscious mice, we used a FosTRAP Cre-dependent tdTomato reporter model to identify activated neurons^36^. Lithium increased the number of TRAP-positive neurons in nodose ganglia, AP and nucleus tractus solitarius (NTS) (Figs 2a, b), consistent with activation of a vagal-brainstem pathway.

**Fig. 2.**
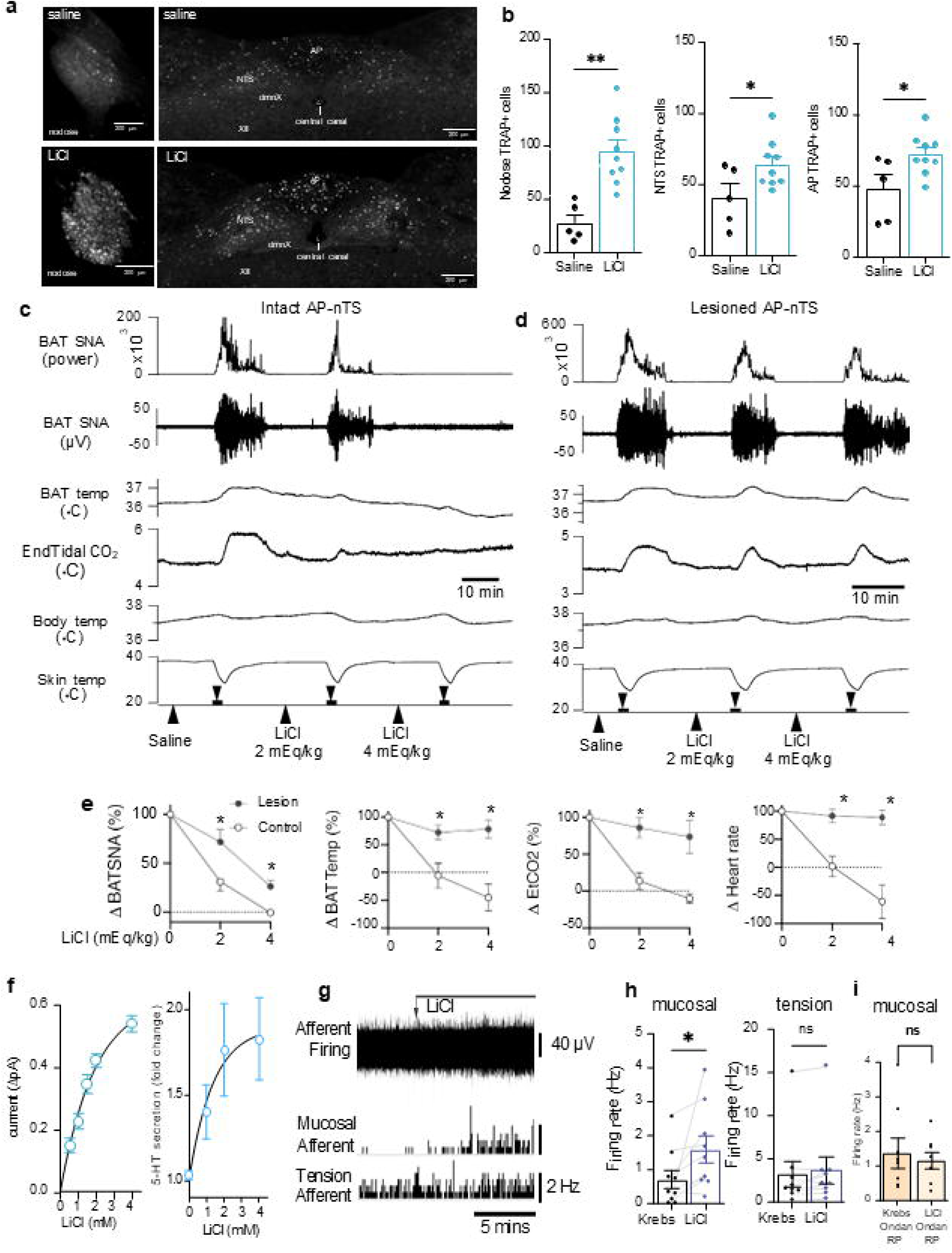
A gut-brain pathway is activated by lithium. **a** Micrographs from FosTRAP+ mice in the nodose ganglion and AP-NTS after i.p. saline or 2mEq/kg LiCl. **b** Group data (saline, n= 5; LiCl n=9, unpaired t-test for nodose f, AP and NTS. **c** In anesthetized paralyzed rats, LiCl inhibits cold-induced increases in sympathetic nerve activity (SNA) to brown adipose tissue (BAT), BAT temperature and end-tidal CO_2_ in intact animals, but **(d)** not in animals with lesions of AP-NTS. Horizontal bars indicate periods when truncal skin was cooled via an ice-water jacket. Upward pointing arrows indicate time of administration of either saline or LiCl (2 and 4 mEq/kg i.p.). **e** Group data for the effect of LiCl on metabolic parameters and heart rate in intact versus AP-NTS lesioned rats (n=6 in each condition, factorial ANOVA and Tukey’s post-hoc correction, *p<0.05). The cold-induced increase in parameters before saline injection was taken as 100%. **f** Dose-dependent increases in 5-HT secretion in mouse duodenum in response to the application of LiCl (1mM) as measured using either electrochemistry (n=13, one way ANOVA and Tukey’s post-hoc correction, non-linear regression, R^2^=0.72) or 5-HT ELISA (n=6, one way ANOVA and Tukey’s post-hoc correction, non-linear regression, R^2^=0.35). **g** Firing in a vagal trunk (top) in response to LiCl (1 mM) and subsequent firing rates from single afferent fibers identified as a mucosal afferent and a tension afferent. **h** Average discharge of mucosal and tension afferent nerves in response to LiCl (mucosal n= 8, tension n= 9, paired t-test). **i** LiCl induced mucosal afferent discharge is prevented by combined 5-HT_3_ receptor antagonist ondansetron (ondan) and NK_1_ receptor antagonist RP67580 (RP) (n=8, paired t-test).

We therefore tested whether this pathway is required to mediate lithium’s thermoregulatory actions. Cold stimulation of the trunk normally evokes robust sympathetic nerve activity to brown adipose tissue (BAT), driving thermogenesis and increasing whole-body metabolic rate. Intraperitoneally administered lithium dose-dependently suppressed this sympathetic discharge, reduced BAT thermogenesis, and lowered metabolic rate measured as percent CO_2_ in the expired air, and heart rate (Fig 2c). These inhibitory effects were abolished by lesions of the AP and the underlying NTS (Fig 2d, e), confirming the necessity of an intact gut-brain pathway for lithium’s actions.

Conditioned taste aversion, a key lithium-associated behavioural change, has recently been shown to require activation of EC cells^29^, the major source of peripheral 5-HT, with synthesis via tryptophan hydroxylase 1 (Tph1) rather than the neuronal isoform Tph2^21,30–32^. EC cells are activated not only by ingested nutrients, but also by potentially harmful agents including chemotherapeutics such as cisplatin^20,21,33^. Genes encoding the enzymes for the production of 5-HT (Tph1) and the neuropeptide substance P (Tac1) are co-expressed in EC cells (Fig S1)^34^, enabling co-release of 5-HT and substance P which activate 5-HT_3_ and neurokinin-1 (NK1) receptors on gastric vagal afferent fibers or in the AP to induce nausea and nausea-associated behaviours^33,35^. These same receptors drive EC cell-activated conditioned taste aversion^29^.

Ex vivo application of lithium to mouse duodenum rapidly and dose-dependently triggers 5-HT release measured either electrochemically or by ELISA (Fig 2f), confirming direct activation of EC cells. We used an *ex vivo* mouse gut preparation to record activity from an extrinsic vagal trunk entering the upper gut. This records activity from all fibres in the trunk, with single units discriminated due to their unique amplitude and duration profiles^22^. Mucosal vagal afferents can be identified by their sensitivity to mucosal stroking, but insensitivity to graded tension (Fig S2). We found that lithium selectively increases discharge of mucosal afferent fibers responsive to gentle stroking of the mucosa, but not of tension-sensitive mechanoreceptors (Figs 2g-h). Combined blockade of 5-HT_3_ and NK1 receptors abolishes lithium activation of mucosal vagal afferents (Fig. 2i), demonstrating that lithium activates vagal afferent terminals via co-release of 5-HT and substance P from EC cells.

## EC cells are essential for lithium action

To investigate whether gut EC cell activation mediates lithium’s physiological and behavioural effects in conscious animals, we employed a Tph1^CreERT^^2^ mouse line that drives changes only in cells expressing Tph1, without affecting Tph1 expression itself. From this we created a Tph1^CreERT^^2/+^;Rosa26^DTA/+^ mouse model^37^ that selectively destroys EC cells after 5 days tamoxifen treatment followed by 5 days washout (Fig 3a-b). We employed a reporter mouse line (Tph1^CreERT^^2/+^;Rosa26^tdTom/+^) to confirm that this approach targets EC cells in the gut lining (Fig S3a-c) but does not affect enteric or brain 5-HT synthesis which is driven by Tph2, a separate enzyme (Fig S3d). Thus, brain neurons associated with 5-HT-synthesis, including those in the dorsal raphe^30–32^ are unaffected in our Tph1^CreERT^^2/+^;Rosa26^DTA/+^ mouse model.

**Fig. 3.**
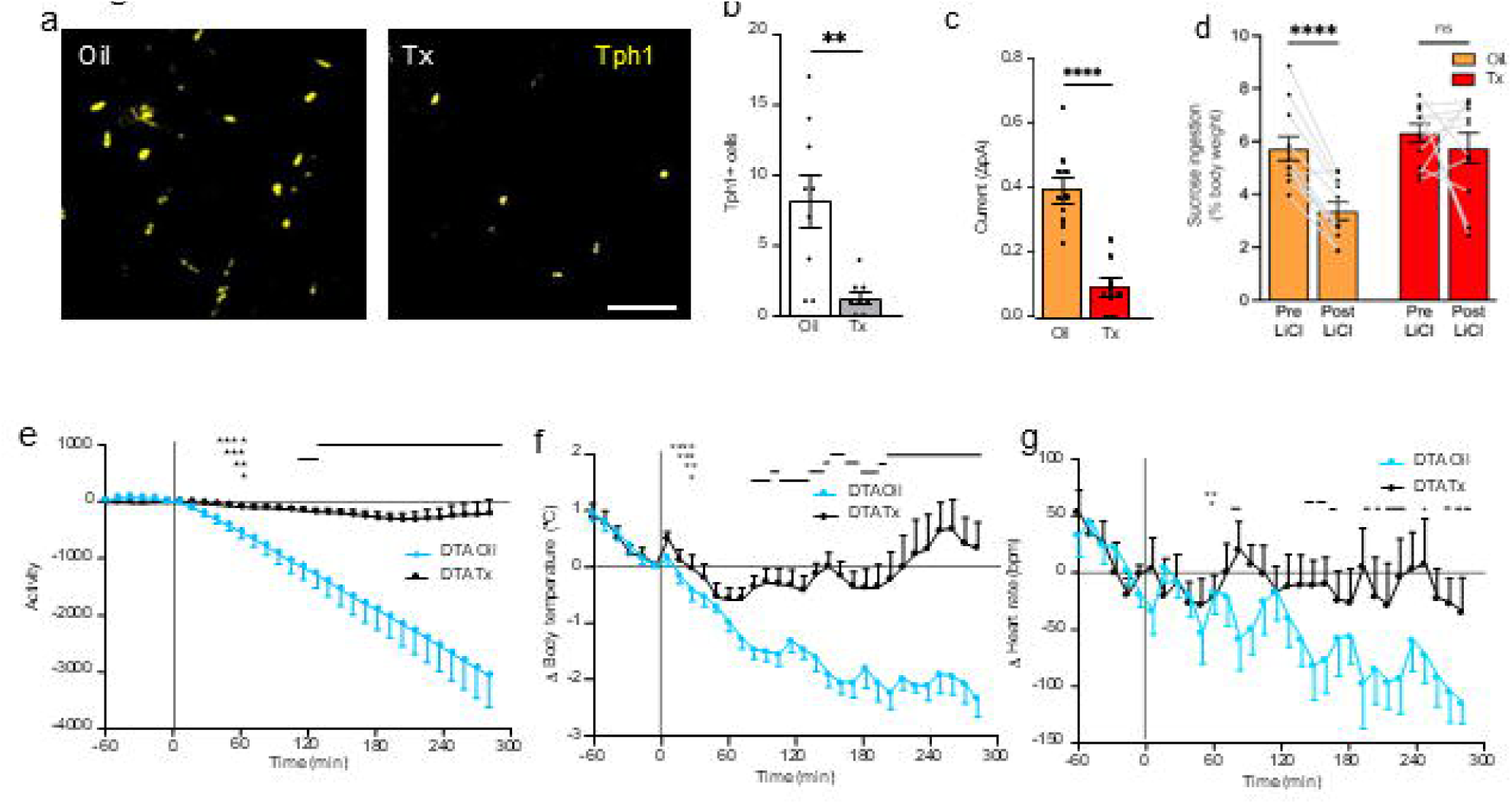
Lithium acts via gut enterochromaffin cells. **a** Tph1+ cells in small intestine of *Tph1^CreERT^*^2^*^/+^;Rosa26^DTA/+^* (*Tph1-DTA*) mice injected with 0.05 mg/kg body weight tamoxifen (Tx) for 5 days or oil control (scale bar = 100_μ_m). **b** Quantification of this data (n= 9, oil; n=8, Tx, unpaired t-test). **c** Electrochemical current change in mouse duodenum in response to LiCl (1 mM) (n=10, oil; n=9, Tx, un-paired t-test). **d** Lithium induced CTA occurs in Tph1-DTA mice treated with oil [paired t-test], but not Tx (n=11, paired t-test). **e-g** Loss of EC cells blocked LiCl-induced changes in locomotor activity, body temperature, and heart rate in freely moving mice (n=12 per group). Changes were analyzed using repeated measures two-way ANOVA, showing significant effects for treatment-time interaction for all three parameters (P<0.0001). Post hoc test identified significant differences at specific time points (marked by black bars with asterisks). Data are mean ± SEM. *p<0.05, **p< 0.01, ***p< 0.001,****p<0.0001.

Lithium-induced 5-HT release from upper gut no longer occurs in the Tph1^CreERT^^2/+^;Rosa26^DTA/+^ mouse model of EC cell ablation (Fig 3c). Remarkably, in conscious unrestrained mice, EC cell ablation completely prevents lithium-induced conditioned taste aversion (Fig 3d). EC cell-ablation also prevents lithium-induced reductions in behavioural activity (Fig 3e), body temperature (Fig 3f) and heat rate (Fig 3g). Even very high lithium doses that cause marked hypothermia and immobility in control mice (Fig S4, Movie 2) fail to do so when EC cells have been deleted (Fig S4, Movie 3).

## Enterochromaffin cell Trpm2 channels contribute to lithium’s action

We hypothesized that some specific aspect of EC cell function must make them receptive to lithium. Because ion channels from the Trp (transient receptor protein) family have been associated with lithium action and bipolar disorder^38,39^, and as some of these channels have been linked to thermoregulation, we examined their expression in EC cells and identified Trpm2 as being the only one of these channels with enriched expression (Fig 4a), ∼250-fold enriched compared to surrounding non-EC cells^20^. This aligns with single-cell analysis of gene expression across all gut epithelial cell types^40^ which confirms Tph1-containing EC cells as the only site of Trpm2 expression in the gut wall (Fig 4b). Further focusing on enteroendocrine cell types^34^ strengthens this notion, demonstrating Trpm2 presence in a specific subset of EC cells (Fig 4c). This is functionally relevant in the gut as we demonstrate that Trpm2 antagonists block lithium-induced 5-HT release *ex vivo* from mouse duodenum (Fig 4d).

**Fig. 4.**
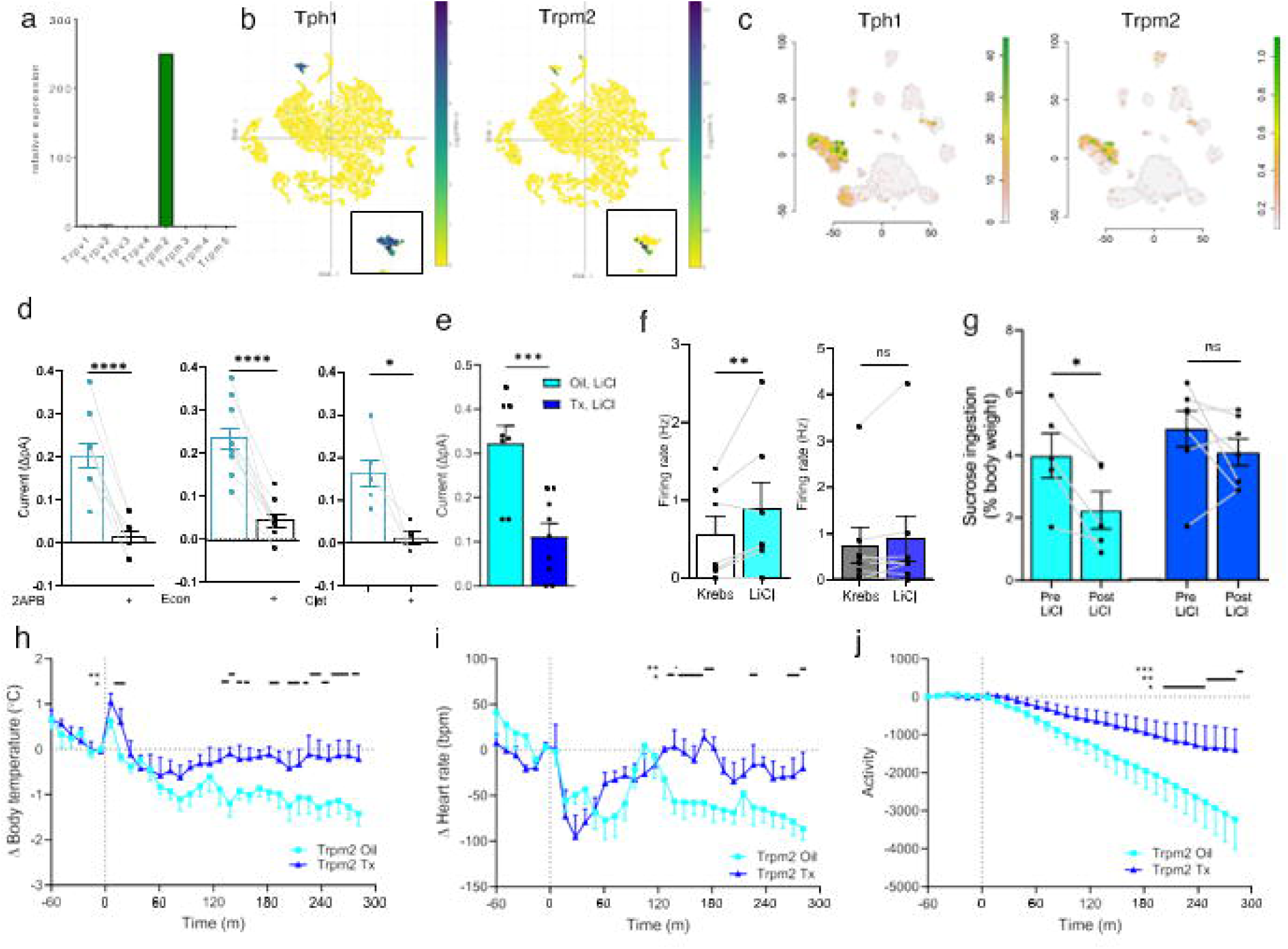
Enterochromaffin cell Trpm2 is essential for lithium’s action. **a** Expression of genes encoding for Trp channels in EC cells. **b** t-distributed stochastic neighbor embedding (t-SNE) plots of single cell expression surveys across all upper gut epithelial cell types in mice showsTrpm2 exists only in a subset of EC cells. **c** Similar gene expression analysis focused on enteroendocrine cells from mouse upper gut also demonstrates Trpm2 expression is highly enriched in a subset of Tph1-containing EC cells. **d** Trpm2 antagonists 2-aminoethoxydiphenyl borate (2APB, 1mM, n= 10), econazole nitrate (Econ, 3_μ_M, n = 11) and clotrimazole (clot, 30 _μ_M, n = 6) reduce LiCl (1 mM) induced 5-HT release in mouse duodenum (paired t-test). **e** LiCl (1 mM) induced 5-HT secretion is reduced in Tx-treated in Tph1^CreERT^^2/+^*;Trpm2^flox/flox^* (Tph1-Trpm2; Trpm2 is ablated from Tph1+ cells) compared to oil-treated controls (n=8 oil; n=9 Tx, unpaired t-test).**f** LiCl (1 mM) increases the firing rate of individual mucosal vagal afferent fibers from oil-treatedTph1-Trpm2 mice (left, n=7) but not from Tx-treated Tph1-Trpm2 mice (right, n=8, paired t-test). **g** Lithium induced CTA occurs in oil-treated Tph1-Trpm2 mice (left, n=5, paired t-test) but not Tx-treated Tph1-Trpm2 mice (right, n=7, paired t-test). **h-j** The effects of LiCl (2 mEq/kg) on body temperature, heart rate, and activity observed in oil-treated Tph1-Trpm2 mice are abolished in Tx-treated Tph1-Trpm2 mice. Body temperature, heart rate, and behavioral activity changes were analyzed using repeated measures two-way ANOVA, showing significant effects for treatment-time interaction (P<0.0001) for all three parameters. Post hoc test identified significant differences at specific time points (marked by black bars with asterisks). Data are mean ± SEM. * p<0.05, **p< 0.01, ***p< 0.001, ****p<0.0001.

To test the importance of EC cell Trpm2 to the *in vivo* actions of lithium, we created an intersectional mouse model, Tph1^CreERT^^2/+^;Trpm2^lox/lox^ mice (herein named *Tph1-Trpm2* mice), and administered tamoxifen for 5 days to eliminate Trpm2 expression only in cells expressing Tph1. This model is specific to EC cells, as single cell gene expression data from cell types throughout the body shows that detectable Trpm2 and Tph1 co-expression occurs only in EC cells (Fig S5).

In the absence of Trpm2 in EC cells in *Tph1-Trpm2* mice, lithium no longer induces 5-HT secretion (Fig 4e) or mucosal afferent nerve discharge (Fig 4f, S6). Importantly, these mice also fail to develop lithium-induced conditioned taste aversions (Fig 4g). Lithium-induced physiological changes, including reductions in body temperature (Fig 4h), heart rate (Fig 4i), and movement (Fig 4j) are substantially reduced in mice lacking Trpm2 expression in EC cells. Thus, EC cell Trpm2 channels are essential mediators of lithium’s actions.

## Lithium releases 5-HT in human gut tissue

To assess the potential translation of our findings to humans, we examined lithium’s effects on surgically removed human gut tissue. Lithium treatment in *ex vivo* human small intestine induces 5-HT release (Fig 5a) and calcium influx in isolated human EC cells (Fig 5b). Single cell analysis of gene expression in human gut epithelial cells revealed *Trpm2* expression to be nearly exclusive to EC cells (Fig 5c). Pre-treatment with Trpm2 antagonists attenuated lithium’s effects in both human gut tissue (Fig 5d) and isolated human EC cells (Fig 5e). Thus, the lithium detecting apparatus in mouse gut is also functional in human gut.

**Fig. 5.**
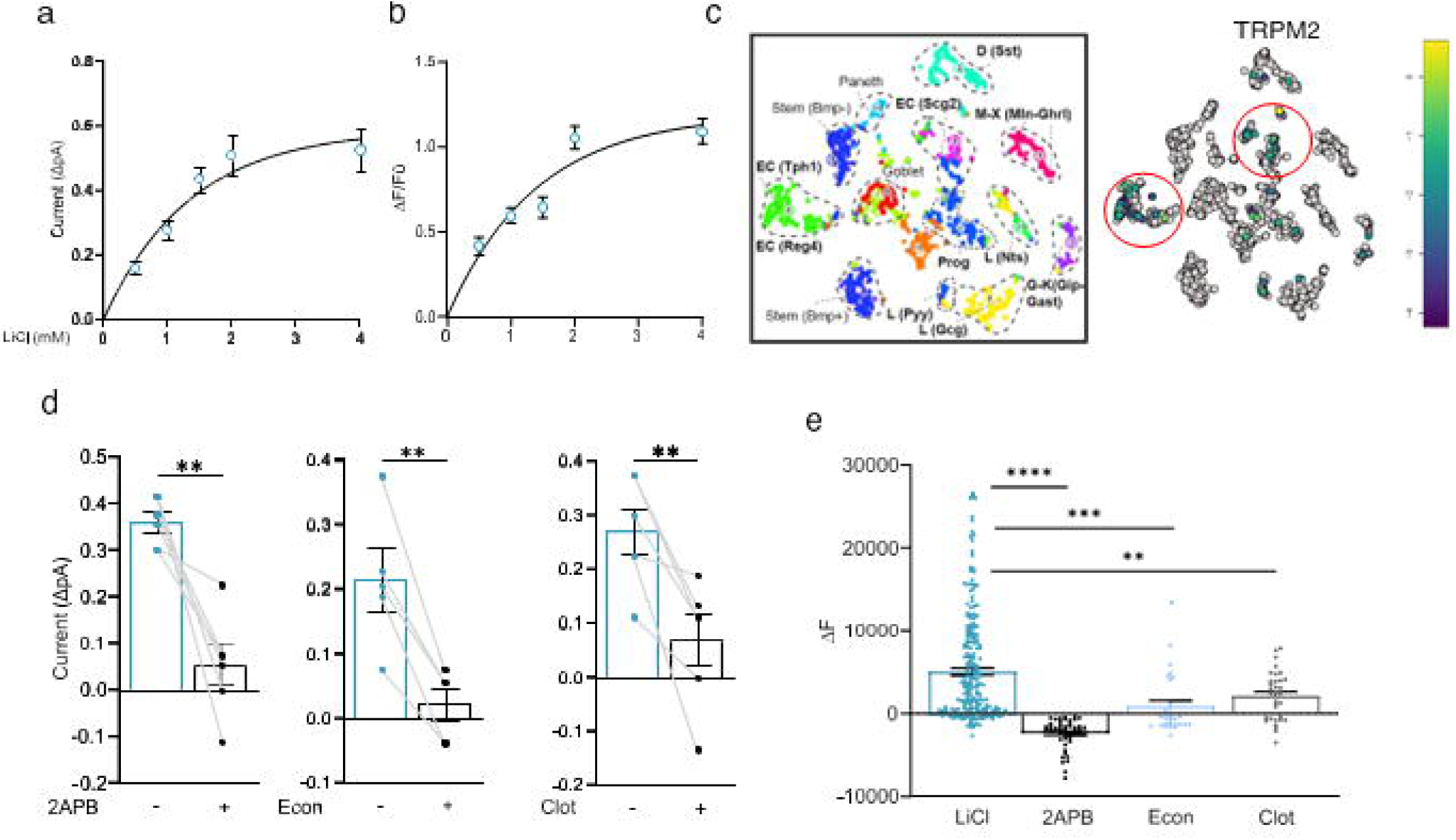
Lithium action in human gut. **a** 5-HT secretion in human ileum increases in response to increasing concentration of LiCl (n = 21). **b** Intracellular calcium in human EC cells increases with increasing LiCl concentrations (n = 722 cells). **c** t-SNE plot displaying the human enteroendocrine cell atlas, each color identifying 13 separate enteroendocrine cell clusters (left) and map displaying the major areas of expression of TRPM2 (red circles) in the different human enteroendocrine cell subtypes from intestinal organoids. Bar displays color-coded unique transcript expression (logarithmic scale). **d** 5-HT secretion induced by LiCl (1 mM) in human ileum is reduced in the presence of the TRPM2 antagonists 2-aminoethoxydiphenyl borate (2-APB, 1mM, n = 6), econazole nitrate (Econ, 3_μ_M, n = 5) and clotrimazole (Clot, 30 _μ_M, n = 6) [paired t-test]. **e** Changes in calcium indicator fluorescence in response to LiCl (1 mM, n = 221 cells) is blocked by the TRPM2 antagonists 2APB (n = 50 cells), Econ (n = 29 cells) and Clot (n = 33 cells). [unpaired t-test]. Data are mean ± SEM. * p<0.05, **p< 0.01, ****p<0.0001, NS, not significant.

### Discussion

Lithium’s therapeutic mechanism remains unknown despite 75 years of successful use for treating the manic phase of bipolar disorder. Traditionally, the primary investigative target has been the brain itself, with suggested mechanisms common to all neurons and thus unable to explain lithium’s specific anti-manic action. We demonstrate, across multiple animal species, and in human gut, that lithium’s primary site of action lies not in the brain but in the gut-brain axis, specifically on gut EC cells that release 5-HT, substance P and other agents. Patients treated with lithium have elevated plasma levels of 5-HT, ^41^ suggesting that lithium activates EC cells to release 5-HT in humans as we demonstrate in our *in vitro* preparations of human intestine.

Lithium directly activates EC cells via their unique expression of Trpm2 ion channels within the gut wall. Adult mice lacking Trpm2 expression in EC cells fail to release 5-HT in response to lithium, or to activate mucosal afferent nerves, or to exhibit lithium-induced behavioural and physiological changes. The magnitude of the effect on blocking lithium’s *in vivo* actions by Trpm2 ablation resembles that seen in the EC cell ablation model. This is unsurprising given that we also demonstrate that the only site in which Trpm2 is expressed in Tph1-containing cells is in gut EC cells. EC cell Trpm2 therefore appears essential for lithium’s physiological and behavioural actions. Lithium activation of EC cells tissue also occurs via a Trpm2-dependent process in human gut, providing a clinical link between our findings in mice and the mechanism of action for lithium’s mood-stabilizing effects in humans.

The peripheral pool of Tph1-synthesized 5-HT does not cross the blood-brain-barrier, so that effects on brain function are mediated via a paracrine action on vagal afferent fibres and via circumventricular organs, including the AP. Lithium engages specific brain pathways precisely because its activation of EC cells sends specific information to the lower brainstem. Peripherally circulating 5-HT is already known to have arousal-reducing actions. Infusion of 5-HT into the arterial supply of the AP causes slowing of the EEG^42,43^, a marker of reduced arousal, similar to lithium’s EEG effects in human patients^44^. We suggest that reduction of arousal, body temperature, metabolic rate and heart rate by lithium is a brain-integrated physiological response that protects the body from potentially toxic agents. Reduced psychological, behavioural and autonomic arousal, with reduced body temperature, metabolic rate and heart rate occur during torpor, also a protective behavioural and physiological “withdrawal” state controlled by specific brain centres^45^, and lithium’s primary gut action provides specific access to these centres.

NTS and AP neurons project to the pontine parabrachial nucleus (PBN) which functions as the “house alarm” for visceral signaling^46,47^. Lithium specifically activates PBN calcitonin gene related peptide neurons, and their optogenetic activation induces conditioned taste aversions^48^. These neurons project directly to the amygdala and the insular cortex^49^ which relay to the medial frontal cortex (in humans, the anterior cingulate portion of the orbitofrontal cortex), forebrain substrates associated with arousal and our interoceptive self-awareness^50^. Thus, the specific brain pathways activated via lithium’s effect on the gut-brain axis are precisely those associated with emotional regulation^51^. Induction of conditioned taste aversion by lithium causes long term rewiring of brain neural circuitry, perhaps highly relevant to lithium’s long term therapeutic actions^52^.

While mood cannot be directly measured in rodents, lithium’s behavioural and physiological effects in animals closely mirror its therapeutic effects in bipolar disorder. Lithium dose-dependently induces conditioned taste aversions in rats well within lithium’s therapeutic window (0.5 to 1.2 mEq/kg)^6^, similar to our demonstration of lithium inducing secretion of 5-HT from EC cells and activating vagal afferent nerves. Lithium substantially reduces exploratory/aggressive responses to an intruder animal at 1 mEq/kg, also within the therapeutic range, just as it reduces aggression in humans^26,27^. The resident intruder model is used to assess aggression in rodents, a symptom sometimes associated with the manic pole of bipolar disorder^53^. Thus, our data link the physiological and behavioural changes induced by lithium via EC cell Trpm2 to its therapeutic actions in bipolar disorder.

Our novel gut-brain axis framework for lithium’s actions (Fig. 6) offers a comprehensive explanation for both its therapeutic efficacy and its gastrointestinal side effects. By aligning behavioural and physiological endpoints with lithium’s known effects in humans, we establish a direct translational framework to investigate its mechanism of action via the gut-brain axis. We suggest that mood stabilization can be achieved by activating specific components of the gut-brain axis, providing novel targets for new therapies for mood disorders.

**Fig 6.**
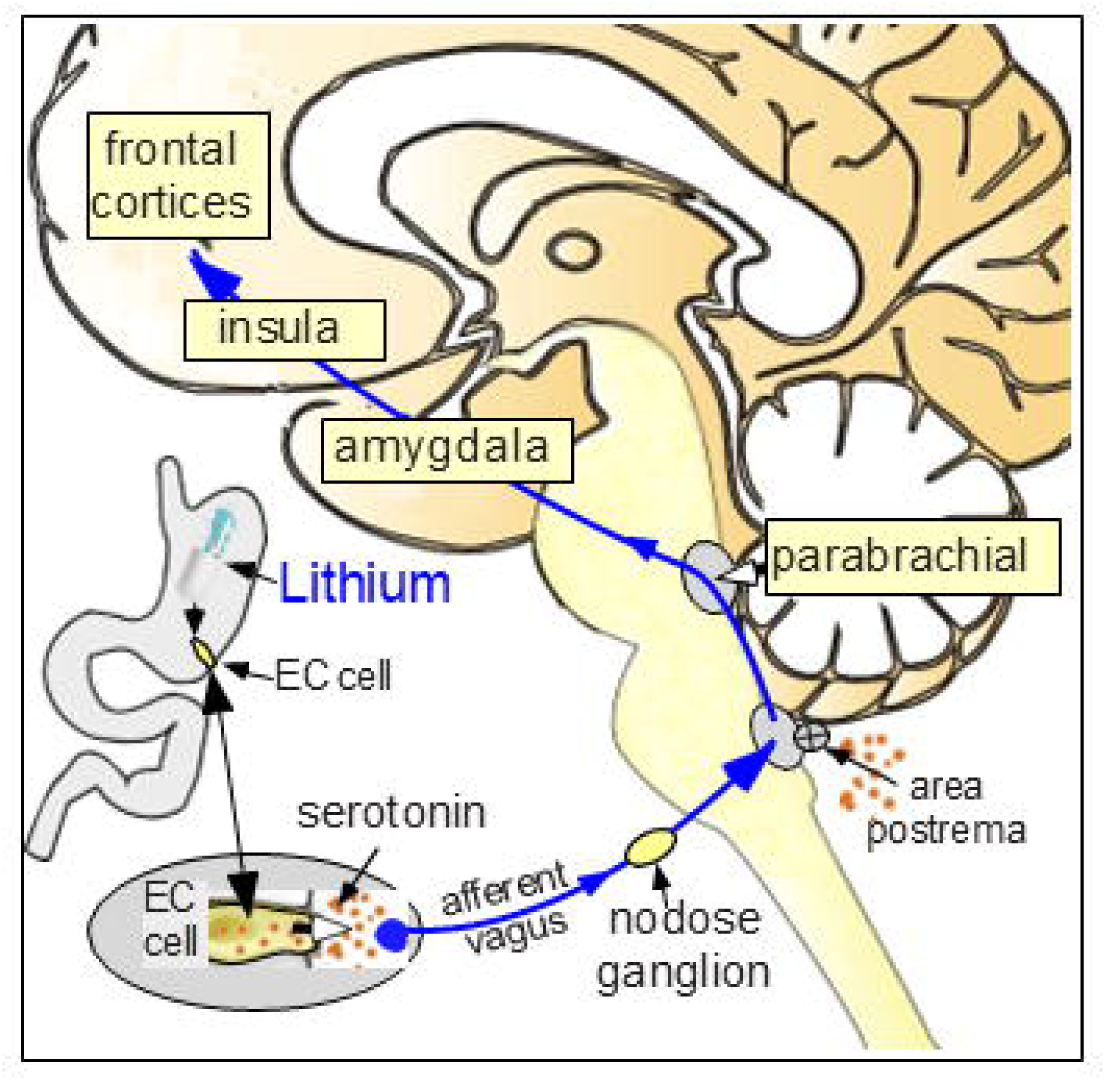
Gut-brain axis pathway for lithium’s actions. Diagram summarizing our proposed neural pathways whereby LiCl reduces behavioral and physiological arousal.

## Acknowledgments

The authors thank the staff of the Department of Surgery of Flinders Medical Centre for their assistance in obtaining surgical specimens. This includes Paul Hollington, Luigi Sposato, Dayan DeFontgalland, Phillipa Rabbit, Tiong Cheng Sia, Abdullah Rana and Chris Liyanage. The authors also thank Rishun Sakai, Zenxi Wang and Kyogo Sakai for technical assistance. The authors acknowledge the facilities, and the scientific and technical assistance of Microscopy Australia (ROR: 042mm0k03) enabled by NCRIS and the government of South Australia at Flinders Microscopy and Microanalysis (ROR: 04z91ja70), Flinders University (ROR: 01kpzv902).

## Funding

This work was supported by the National Health and Medical Research Council and the Baszucki Brain Research Fund.

## Author contributions

Conceptualization: DJK, WWB.

Investigation: all authors.

Funding acquisition: DJK.

Writing – original draft: DJK, WWB, DWT, YO, RAP.

Writing – review & editing: all authors

## Supplementary Materials

Materials and Methods

Supplementary Text

Figs.S1 to S10

Tables S1 to S2

References (*44*–*56*)

Movies S1 to S3

## Materials and Methods

### Animals

All care and procedures involving animals were carried out in accordance with protocols approved by the Animal Ethics Committees of Flinders University (Mice: #5147-19, Rats: #691-18 and Guinea Pigs: #940-17) and the Monell Chemical Senses Center (#1190). Mice (strain specific details below), Sprague-Dawley rats, and Tri-colour guinea pigs (males) were used. Animals were housed in a temperature-controlled, pathogen-free environment, with a 12-hour light/dark cycle and *ad libitum* access to food and water except during surgical procedures and/or behavioral tests. Guinea pigs were housed individually, with a “companion” guinea pig in an adjoining cage.

#### Mice

6–12-week-old male and female mice were used in this study, in accordance with protocols approved by the Animal Ethics Committee of Flinders University (ethics no: 5147-19) and the Monell Chemical Senses Center (#1190). The following strains were used: bred in house (Flinders University) C57BL/6J, Tph1^creERT2/+,^ Tph1^creERT2/+^;ROSA26^DTA/+^, Tph1^creERT2/+^;Trpm2^lox/lox^ and cFOS^TRAP^ (bred in-house (Monell Chemical Senses Center) from FosTRAP Jax B6.129(Cg) Fostm1.1(cre/ERT2)Luo/J (JAX stock no.021882) crossed to Ai9 mice (bred in-house (Monell Chemical Senses Center) from Jax (stock no. 007905). A 5 consecutive day tamoxifen injection protocol (Tx; Sigma-Aldrich; 1 mg/20 g body weight, solubilized in corn oil, administered intraperitoneally) was used to ablate Tph1-expressing cells (EC cells) in Tph1^creERT^^2/+^;ROSA26^DTA/+^ mice and to ablate Trpm2 from Tph1-expressing cells in Tph1^creERT^^2/+^;Trpm2^lox/lox^ mice. Corn oil (Sigma-Aldrich) injected mice were used as controls.

### Human Intestinal Tissue

Small intestinal tissue was obtained from patients undergoing right hemicolectomies for cancer. All patients provided prior written informed consent for tissue donation at either Flinders Medical Centre or Flinders Private Hospital, as approved by the Southern Adelaide Clinical Human Research Ethics Committee (ethics no: 50.07). Samples were taken from the resected margin, as far from the lesioned area as possible. Tissue was transported to the laboratory in room-temperature Krebs solution (NaCl 118mM, KCl 4.8mM, CaCl_2_ 2.5mM, MgSO_4_ 1.2mM, NaHCO_3_ 25mM, NaH_2_PO_4_ 1.0mM, glucose 11mM, bubbled with 95% O_2_, 5% CO_2_, pH 7.4) within 15 minutes of resection.

### Guinea Pig Movement studies

#### Radiotelemetry Temperature Transmitter Implantation

Tricolour guinea pigs (male, 385-789g) were placed under general anaesthesia (2-3% isoflurane in 100%). Before implantation, animals received one subcutaneous (s.c.) dose of analgesics (4 mg/kg carprofen [Carprieve, Norbrook, Tullamarine, VIC, Australia] + 0.05 mg/kg buprenorphine [ilium TemVet, Troy Laboratories, Glendenning, NSW, Australia]) and antibiotics (10 mg/kg enrofloxacin [Baytril, Bayer, Pymble, NSW, Australia]). A telemetric temperature transmitter (TA-F40, Data Sciences International [DSI], Transoma Medical, Arden Hills, MN, USA) was then implanted intraperitoneally (i.p.). Once emerged from anaesthesia, the animals were returned to the animal facility for at least one week’s recovery period. For the first two days of recovery, animals received one s.c. injection of carprofen (2 mg/kg) daily.

#### Recording

Animals were placed inside an acrylic open-top cage (350 x 400 x 450 mm) contained within an isolated climate-controlled chamber (0.32 m^3^; Biomedical Engineering, Flinders Medical Centre, Bedford Park, SA, Australia) that was maintained around 22°C with a diurnal light cycle (7am light, 7pm dark). The open-top cage was divided into two separate chambers via a clear, perforated divider. This divider allowed each experimental guinea pig to have a non-experimental companion, thereby reducing solitude induced stress. Body temperature from the telemetry probe were detected with a PhysioTel Receiver (RPC-1, DSI) and a communication box (Matrix 2.0, DSI) and were captured by LabChart (ADInstruments, NSW, Australia) at 10Hz via an analogue-digital converter (Powerlab, ADInstruments). Locomotor activity was recorded with a USB video camera module ((1280×720 30fps wide angle, fisheye lens) ELP, Amazon Japan, Japan) in Labchart (ADInstruments, NSW, Australia), and quantified by measuring total distance moved with EthoVision XT software (Noldus, Wageningen, Netherlands). Mean total distance moved was calculated for each 10-minute period. Locomotor movement trace maps were made with Deeplabcut^54^. The video was manually reviewed to determine the number of seconds per minute spent in “cooling posture”, defined as lying flat on belly or side with at least one leg splayed out.

#### Lithium Dose-Response Protocol

After habituating to the recording chamber for at least 24 hours to stabilize body temperature and behavior, experimental guinea pigs were injected i.p. (left lower quadrant) with a dose of lithium (0.1, 1, 2, 4, 6 mEq/kg LiCl [Sigma-Aldrich]). A 3-day clearance period was ensured before the next dose. The first dose administered to each successive animal was given in alternating order to account for serial effects.

#### Data Analysis

Delta body temperature values were calculated by subtracting the mean of 5– 10 mins before injection from each datapoint. During body temperature recordings, if artifacts due to noise occurred, these were accounted for by interpolating an appropriate value using data points at the surrounding time points. For lethargic behaviour, mean time spent in cooling posture was taken over the first 60 minutes post-lithium; for locomotor activity, we took cumulative distance moved at 240 mins post-lithium. For the lithium dose-response data, linear regression was run on each of these parameters against log-transformed lithium dose. Multiple linear regressions were run between log-transformed dose and cumulative distance moved at a single point in time every 10 minutes after lithium-injection. All data are presented as mean ± SEM.

### Tail and forepaw skin temperature studies

To measure thermoregulatory skin temperature, infrared images of either tail skin (rats) or forelimb skin (guinea pigs) were taken with an infrared camera (A40 FLIR, Wilsonville, Oregon, United States). The maximum skin temperature along the tail or the forelimb was recorded. Guinea pigs were handheld to expose forelimbs when images were taken. One hour after starting infrared camera recordings, animals were injected intraperitoneally with either vehicle (0.9% saline, 0.5 mL) or lithium (2 mEq/kg LiCl in 0.5 mL water for injection). Three days separated each injection. Injections were administered in an alternating order to account for serial effects.

### Brown adipose tissue sympathetic nerve studies in anaesthetized rats

Preparatory surgery was performed as previously described ^55^. Briefly, Sprague-Dawley rats were anesthetized with isoflurane (2% in oxygen) and an endotracheal tube was inserted via a tracheotomy. The right femoral vein and artery were cannulated for drug administration and arterial pressure measurements, respectively. Rats were then anesthetized with urethane (400-800mg/kg) and α-chloralose (40-80mg/kg) (i.v.) as isoflurane anesthesia was ceased. Rats were mounted onto a stereotaxic frame and the dorsal surface of the medulla oblongata was exposed. The area postrema and the nucleus tractus solitarius were removed by aspiration. Rats were subsequently paralyzed with d-tubocurarine (0.3ml i.v. of a 0.3mg/ml solution in ringer) and artificially ventilated with 100% O_2_ (60-65 cycle/min, volume 1ml/100g body weight/cycle). End tidal CO_2_ (Normocap; Datex, Helsinki, Finland) was directly recorded as per previous studies^55,56^ and maintained at 4–5% by adjusting the ventilation volume. Animals were allowed to recover from paralysis, so that adequate anesthesia could be confirmed by absence of the corneal reflex before paralysis was reestablished. Additional urethane/α-chloralose anesthetics was given if necessary. Body, brown adipose tissue (BAT) and skin temperatures were measured with thermocouples (TC-2000; Sable Systems) positioned in rectum, interscapular BAT and abdominal skin respectively. Body temperature was maintained constant by varying the water temperature in a truncal jacket^57^.

An interscapular BAT sympathetic nerve was isolated^57^. Nerve activity was recorded by a pair of silver electrodes and amplified (gain 20,000, NL104 amplifier, Digitimer Ltd., Welwyn Garden City, U.K.) and filtered (band-pass 1–1000 Hz, NL125 filter, Digitimer). BAT sympathetic nerve activity (BATSNA), evoked by briefly perfusing ice water through the water jacket. Either saline or LiCl was administered (i.v.) before the cooling procedure and effects on BATSNA, BAT temperature and end-tidal CO_2_ were assessed. At the end of experiments, a ganglionic blocker, hexamethonium bromide (Sigma, H0879) (10 mg/kg i.v.) was administered to confirm loss of BATSNA.

#### Data analysis

Data were sampled and digitized by PowerLab at 1 kHz (ADInstruments Inc., Bella Vista, NSW, Australia). Heart rate was computed from electrocardiogram (ECG). Data was analyzed using Igor Pro (WaveMetrics, Portland, OR), and with Graph-pad Prism 6 (Graph-pad software Inc., CA). Signal artifacts were replaced with interpolating values from surrounding time points. Amplitude of BATSNA was expressed as total power spectral (between 0-20Hz, 5.12-s segment)^55^. Cold-induced changes in BATSNA, BAT temperature, end-tidal CO_2_ and heart rate were recorded as the peak value in their twice differentiated wave form at onset and termination of cold-induced changes in the skin temperature. Response amplitudes, calculated as the area under the respective traces between the start and end of truncal cooling before any treatment, were set as 100%.

### TRAP protocol

The TRAP protocol was carried out as reported previously^36^ to label cells responsive to LiCl or saline. Mice were habituated to handling and intraperitoneal (i.p) saline injections for 3 days before the TRAP protocol. Briefly, mice were fasted 6 hours, and either LiCl (2mEq/kg) or isotonic saline (control) was administered i.p. 3 h after “lights-on”. Three hours after the LiCl administration, mice were injected with 4-hydroxytamoxifen (4-OHT; 30 mg/kg, i.p.; Millipore Sigma, Burlington, MA). Food was returned to the home cages 3 h later.

#### Perfusion and tissue collection

Mice were sacrificed 2 weeks after the trapping protocol. Anesthetized animals were transcardially perfused with phosphate buffer saline (PBS), followed by 4% paraformaldehyde (PFA) in PBS. Brains and nodose ganglia were collected, postfixed for 24 h in 4% PFA, and transferred to 30% sucrose with 0.1% sodium azide for cryoprotection. Cryosections (35 µm) of the brainstem were obtained for imaging of the dorsal vagal complex (DVC) using a Leica microtome (CM 3050 S, Leica Biosystems) at -20 °C. Sections corresponding to the DVC were mounted on Fisherbrand Superfrost Plus slides (Fisher Scientific) and cover-slipped with Prolong Diamond Antifade Mountant (Invitrogen, Waltham, MA) for imaging. Whole nodose ganglia were mounted and cover-slipped immediately before imaging.

#### Imaging

Brainstem tissue sections corresponding to the DVC were imaged using a Keyence BZ-X800 in a single plane with autofocus capture at 10x. Nodose ganglia were imaged with a 2-Photon microscope at 16x (Bruker), and 10um Z-stacks of the whole nodose were obtained. Quantification of the Td-tomato-positive cells was performed on ImageJ software after adjusting brightness and contrast using the “analyze particles” function. For the DVC, the count of positive cells was averaged from three representative sections (corresponding approximately to -7.76,-7.56, -7.32, from bregma according to^58^. For the nodose, the total amount of positive cells was counted from the max projection of Z-stack and the sum of both nodose ganglia per animal was reported.

#### Data Analysis

Data are presented as Mean ± SEM and were analyzed using a Student’s t-test in the software GraphPad Prism 9. Statistical significance was established at p<0.05.

### Resident-Intruder rat studies

#### Radiotelemetry Temperature Transmitter Implantation

All preparatory surgical procedures were performed under general anaesthesia (2% isoflurane (Veterinary Companies of Australia, Kings Park, NSW, Australia) in 100% oxygen). Before implantation, animals received one subcutaneous (s.c.) dose of analgesics (0.02 mg/kg buprenorphine) and antibiotics (10 mg/kg enrofloxacin, Baytril). A telemetric temperature transmitter (F40-TT, Data Sciences International [DSI], Transoma Medical, Arden Hills, MN, USA) was then implanted intraperitoneally (i.p.). Following surgery, analgesia (meloxicam 1mg/kg s.c., Troy laboratories, New Zealand) were administered. Once the rat had recovered, it was individually caged and returned to the animal holding room for at least one-week. Once all experimental recordings were completed, the animal was euthanized with pentobarbitone sodium (180 mg/kg i.p.) (Virbac Pty Limited, Milperra, NSW, Australia).

#### Recording of physiological parameters, drug administration and the intruder-stress model

The day before of experiments, the resident rat was placed into a plastic, open roofed, ‘home cage’ (350 mm wide x 400 mm long x 450 mm height) located in a temperature-controlled recording chamber (Biomedical Engineering, Flinders University). The ambient temperature in the chamber was maintained at 22^°^C. On the of experiments, either vehicle (saline, s.c.) or LiCl (1, 2, or 4mEq/kg s.c.) was administrated. Thirty minutes after the administration, the chamber was opened, and a second male rat (intruder rat) confined to a small plastic and wire mesh cage (19 x 29 x 12 cm) was introduced to the home cage. Recording of the experimental parameters continued throughout the 30-min intrusion period. Drugs were administered in a counterbalanced rotating order to control for serial effects, with three days between each injection^59,60^. Locomotor activity was detected with a passive infrared sensor, and digitized at 10Hz (NaPiOn, AMN1111, Panasonic, Osaka, Japan).

#### Data analysis

Physiological signals were recorded with PowerLab and analyzed with Igor Pro. During body temperature recordings, if artifacts due to noise occurred, these were accounted for by interpolating an appropriate value using data points at the surrounding time points. To evaluate the effect of LiCl or saline on intruder-elicited changes in body temperature, the difference between the mean of the 10 min period prior to drug administration and the mean of the 30-min intrusion period was calculated (delta body temperatures from the pre-injection level). For locomotor behavioral activity, signals from the infrared activity sensor were cumulatively added from 30 minutes prior to drug administration. The cumulative value at the time of drug administration was set to 0. The cumulative value at the time when the intruder was removed was used for analysis. Linear regression was run on each of these parameters against log-transformed lithium dose for dose-response analysis. Group results (mean ± SEM) were calculated for each experimental condition.

### Electrophysiology

#### In vitro mouse gastric-vagal afferent preparation

C57BL/6J and Tph1^CreERT^^2/+^; Trpm2 ^lox/lox^ mice were used for vagal afferent recordings. Mice were euthanised by isoflurane inhalation and cervical dislocation. The stomach and the oesophagus with attached vagal innervation were excised and placed in a petri dish lined with Sylgard (Dow Corning, Midland, MI, USA) containing Krebs solution (in mM: NaCl, 118; KCl, 4.75; NaH_2_PO_4_, 1.0; NaHCO_3_, 25; MgCl_2_, 1.2; CaCl_2_, 2.5; glucose, 11; bubbled with 95% O_2_/5% CO_2_). Vagal innervation was exposed along the esophagus to where individual nerve trunks entered the stomach; then the esophagus was removed. The stomach was opened along the greater curvature creating a flat sheet preparation, mucosa facing up. Several nerve trunks (2–4 trunks) entering the preparation were mobilized for 5–8 mm. Preparations were transferred to a 10 ml, Sylgard-lined recording chamber, and the dissected nerve trunks were pinned, under slight tension, to the Sylgard, using 50 _μ_m tungsten pins. The chamber was continuously superfused with warmed, oxygenated Krebs solution (37°C). A paraffin oil bubble was attached to the base of the chamber, and the end of the dissected nerve trunk was then pinned inside the oil bubble. Recordings were made from the nerve with a 100 _μ_m Pt/Ir wire electrode insulated apart from the last 400 _μ_m. Signals were amplified (ISO-80, WPI, Sarasota, FL, USA), recorded at 20–40 kHz (PowerLab16s/p, LabChart 7 Pro software, AD Instruments, Sydney, NSW, Australia). Single units were discriminated by amplitude and duration using Spike Histogram software (AD Instruments, Sydney, NSW, Australia).

#### Classification of mucosal vagal afferents

Mucosal afferents were identified by their sensitivity to mucosal stroking, but insensitivity to graded tension. With the stomach under minimal resting load (∼0.5g) mechano-transduction sites were identified by stroking the mucosal surface with calibrated von Frey hairs (40mg, 160mg) to evoke bursts of action potentials. Tension was applied via a purposed built array of hooks attached to one edge of the preparation, via a cotton thread, to an isotonic transducer (52-9511, Harvard Biosciences, South Natick, MA, USA) with (minimal) resting load 0.5g. Stretches were applied by adding counterweights (1-4g) for 30s to the isotonic lever; these constant loads stretched the stomach in a circular direction, while the transducer recorded changes in length. The chemosensitivity of mouse gastric vagal afferents was determined after mechanical properties were established. Ten minutes of normal baseline activity was maintained before the addition of a drug via superfusion and a bolus dose to bring the bath to concentration. LiCl (1mM) in Krebs solution was applied via superfusion for 15 minutes.

#### Antagonist studies

We determined the effect of combined 5HT_3_ receptor (ondansetron, Tocris RDS163550) and NK-1 receptor antagonism (RP67580, Tocris Bioscience, RDS289150) on vagal nerve firing in response to LiCl. Antagonists were applied to the preparation via superfusion for 10 minutes prior to the addition of 1mM LiCl. Ondansetron was dissolved in dH_2_O and diluted in Krebs solution on the day of the experiment to the final concentration of 10^-5^M. RP67580 was dissolved in DMSO and diluted in Krebs solution on the day of the experiment to the final concentration of 10^-8^M.

#### Data analysis

Statistical comparisons were performed using Prism 9 software (GraphPad Software, Inc., San Diego, CA, USA). Two means were compared using paired two-tailed t-tests were used to compare the effects of applied solutions to that of baseline. Differences were considered significant if P< 0.05.

### Amperometry

Segments of duodenum from male and female C57BL/6J, Tph1^creERT^^2/+^;ROSA26^DTA/+^, and Tph1^creERT^^2/+^;Trpm2^lox/lox^ mice of 8–12 weeks of age were excised and placed in Krebs solution. Following the mesenteric border an incision was made along the duodenum and was pinned mucosal side up in a Sylgard lined petri dish containing Krebs solution. Fresh human ileum was also collected, the mucosa dissected and pinned luminal side up in a Sylgard lined petri dish containing Krebs solution. Real time secretion of serotonin was measured with carbon fibre amperometry^61,62^. Oxidation of serotonin was induced through the application of a 400mV current, a carbon fibre probe (ProCFE, Dagan Corporation, Minneapolis, MN) above the tissue and EPC-10 amplifier and Pulse software (HEKA Electronic, Germany) with the current sampled at 10 kHz recorded the change in current from the oxidation of serotonin. Krebs solution was the standard bath solution, all solutions were continuously applied to the tissue using a gravity perfusion system. Experiments with C57BL/6J mouse duodenum and human ileum were conducted with varied concentrations of LiCl (0.5, 1, 1.5, 2, or 4 mM) to obtain a dose response. Antagonist studies were conducted with tissue pre-incubation of the antagonists 2APB (1mM), econazole nitrate (3µM), or clotrimazole (30µM) followed by stimulation with Krebs containing 1mM LiCl and antagonist. Recordings from tissue derived from Tph1^CreERT^^2^;Rosa26^DTA/+^ and Tph1^CreERT^^2/+^;Trpm2^lox/lox^ mice compared serotonin secretion at baseline vs. LiCl (1mM).

#### Data analysis

Recordings and analysis were undertaken in a paired fashion to minimize variance between different tissue preparations or time. Thus, we analysed the effect under control (Krebs) and then under experimental (LiCl) conditions.

### Immunohistochemistry

Duodenum from Tph1^creERT^^2/+^;ROSA26^DTA/+^ mice exposed to a tamoxifen or corn oil injection protocol was pinned under maximal tension and fixed overnight in 4% paraformaldehyde at room temperature. Preparations were permeabilized overnight (20% DMSO, 2% Triton X-100 in PBS), blocked for 1 h (10% normal donkey serum), then incubated with anti-Tph1 antisera (Merck, AB1541; sheep; 1:200) for 2 nights. Labelling was revealed by incubating with donkey anti-sheep Cy™3 overnight (Jackson, 713-165-003; 1:200). Z-stack images were captured at 20x magnification (NA: 0.8) with 2 µm spacing for analysis on a LSM880 confocal microscope (Zeiss, Oberkochen, Deutschland). For examining Tph1 expression in brain and gut, Tph1 cells were genetically labelled with dtTomato in *Tph1^CreERT^*^2^*^/+^;Rosa26^tdTom/+^* (*Tph1-tdTom*) mice at 7 days following tamoxifen injection. Sagittal sections in the brain, jejunum, and colon, were co-labelled with antibodies against Tph1 (Thermofisher, PA1777, 1:100), Tph2 (Thermofisher, PA1778, 1:100), or 5-HT (LifeSpan BioSciences, LS-C75755, 1:200).

### Condition Taste aversion

Tph1^creERT^^2/+^; ROSA26^DTA/+^ (n = 11 Tx and oil) and Tph1^creERT^^2/+^; Trpm2^lox/lox^ (n= 7 Tx and n = 6 oil) mice were administered tamoxifen to ablate Tph1-positive cells or Trpm2 from Tph1-positive cells respectively. Corn oil was used for controls. All mice were naïve to sucrose or Li prior to the CTA. Mice were habituated for one night to the housing and water bottle before the CTA experiment. CTA experiments were performed on 6 consecutive days in housing chambers during the light cycle. Prior to each day of CTA, mice were water deprived overnight (15-17 hr). On days 1,2 and 4,5, mice had access to water for 20 mins in chamber. On day 3, mice were given access to a novel 10% sucrose solution for 20 mins, followed by an injection of Lithium (2mEq/ml). After the injection mice were placed back into the chamber without access to food or water for another hour, after the hour they were given access to food and water for the rest of the day. On day 6, mice were given access again to the novel 10% sucrose solution for 20 mins.

#### Data analysis

We calculated the amount of water or sucrose solution drunk on each day after the 20 min access period. The sucrose intake ratio was also calculated, comparing sucrose intake after one trial of conditioning (day 6) versus the naïve condition (day 3) and normalized to body weight.

### Telemetry

Mice (Tph1^creERT2/+^, Tph1^creERT2/+^;ROSA26^DTA/+^, Tph1^creERT2/+^;Trpm2^lox/lox^ 19-31 g) were anaesthetised using a general anaesthetic (2% isoflurane, Veterinary Companies of Australia, Sydney, Australia) combined with 0.8L/min oxygen. Analgesia (buprenorphine, 0.05mg/kg, s.c. Troy Laboratories Pty. Ltd, NSW, Australia) and antibiotics (Baytril, 10mg/kg, s.c., Bayer Aust, NSW, Australia) were administered prior to surgery. A telemetry probe (ETA-F10, DSI) was surgically implanted in the intraperitoneal cavity of the mice to measure core body temperature and ECG ^63^. The electrical leads for ECG were passed subcutaneously between the connective tissue and skin, with the negative electrical lead in a thoracic muscle and the positive in the right leg muscle. At the end of surgery, meloxicam (5mg/kg, s.c., Troy Laboratories Pty. Ltd, NSW, Australia) was administered. After surgery, mice were housed individually and allowed to have a week’s recovery before experimentation commenced. For the first 3 days after surgery, mice were given water containing analgesia (oral suspension meloxicam, 0.01mg/ml, Dechra Veterinary Products Pty Ltd, NSW, Australia). The cages maintained a 12h/12h light-dark cycle (on at 0700 and off at 1900).

#### Experimental procedures

Prior to experimental recordings, mice underwent the 5-day consecutive tamoxifen injection protocol described above. Animals were given a two-day rest period between the injections and telemetry recordings. On the day of the recording mice were relocated to the experimental room which was maintained at ∼23°C. Mice were individually placed into Perspex recording cages with ad lithium access to food and water. After 2-hours of habituation, LiCl (2mg/kg) was administered intraperitoneally between 10am – 1pm and parameters recorded for another 5 hours.

#### Data recording and analysis

Body temperature and ECG signals from the telemetry probe were detected with a PhysioTel Receiver (RPC-1, DSI) and a communication box (Matrix 2.0, DSI) and were captured by LabChart (ADInstruments, NSW, Australia) at 1kHz via an analogue-digital converter (Powerlab, ADInstruments). Locomotor activity was detected with a passive infrared sensor (NaPiOn, AMN1111, Panasonic, Osaka, Japan) and digitized at 100Hz. Activity was expressed as the total amount of movement detected (sec per min). Data was then imported into IgorPro (WaveMetrics, OR, USA) for further analysis. Data between genotypes were analysed using un-paired t-tests at each time point.

### Bioinformatic analysis of single-cell data

All commands for the reanalysis of the single-cell sequencing are available at (https://gist.github.com/beardymcjohnface/c096ce52e0697a28fbd8cc0a144955d0). Briefly, single cell sequencing data, consisting of raw read counts of all genes and cells, were downloaded from the Gene Expression Omnibus^34,40,64–66^. Transcript counts were converted to counts per Million (CPM). The log2 of the mean CPM and the fraction of cells expressing a gene were calculated for each gene of interest across the different cell types and visualised in R. We identified co-expression of Tph1 and Trpm2 for a range of tissues and cell types by quantifying the mean expression and fraction of Tph1 (CPM > 100), and separately for Tph1 expression in the subset of cells that also express Trpm2 (CPM > 100). Only datasets containing a cell type with greater than 1% of cells expressing Tph1 were included in visualisations.

### Calcium imaging

48 hours before imaging was undertaken 50B11 and QGP-1 cells were plated on 35mm µ-dishes (DKSH, Ibidi, Australia). 50B11 cells were plated at 10×10^4^ cells per ml on dishes coated with Poly-L-Ornithine, in 2 ml Neurobasal media (10% FBS, 200µl glucose, 300µl B27 supplement). QGP-1 cells were plated at 3×10^4^ cells per ml in 2ml RPMI media (10% FBS, 1% Pen Strep). Live cell calcium imaging was undertaken as previously described ^67^. Briefly, QGP-1 and 50B11 cells were treated with Fluro-4 AM (5µM) in Krebs solution for 45 minutes at 37°C. Cells were maintained in Krebs solution and stimulated with Krebs solution containing either 0.5, 1, 1.5, 2, or 4 mM LiCl, or 70 mM KCl. For antagonist studies, QGP-1 cells were incubated for 10min with antagonists 2APB (1mM), Econazole Nitrate (3µM) or Clotrimazole (30µM) then stimulated with Krebs containing 1mM LiCl and antagonist respectively. Calcium influx was visualised as increases in cell fluorescence on a Cascade II fluorescent microscope (Photometrix, USA). Results were analysed on Imaging Workbench software (Version 6.0.22, INDEC BioSystems, Inc, USA).

### High dose lithium exposure

Tph1^creERT^^2/+^;ROSA26^DTA/+^ mice administered either oil or tamoxifen (5 days), and Tph1^creERT^^2/+^ administered tamoxifen as an additional control group, were injected with a high LiCl dose that was 5-10 times greater than that which causes maximal conditioned taste aversion (16.8 mEq/kg)^6^. Mice were monitored closely for 24 hours, and then rectal temperature measured.

## Notes

### Competing Interest Statement

The authors have declared no competing interest.

