## Supplemental Figures for "The mood stabilizer lithium alters behaviour and physiology via the gut brain axis"

### Slide 1
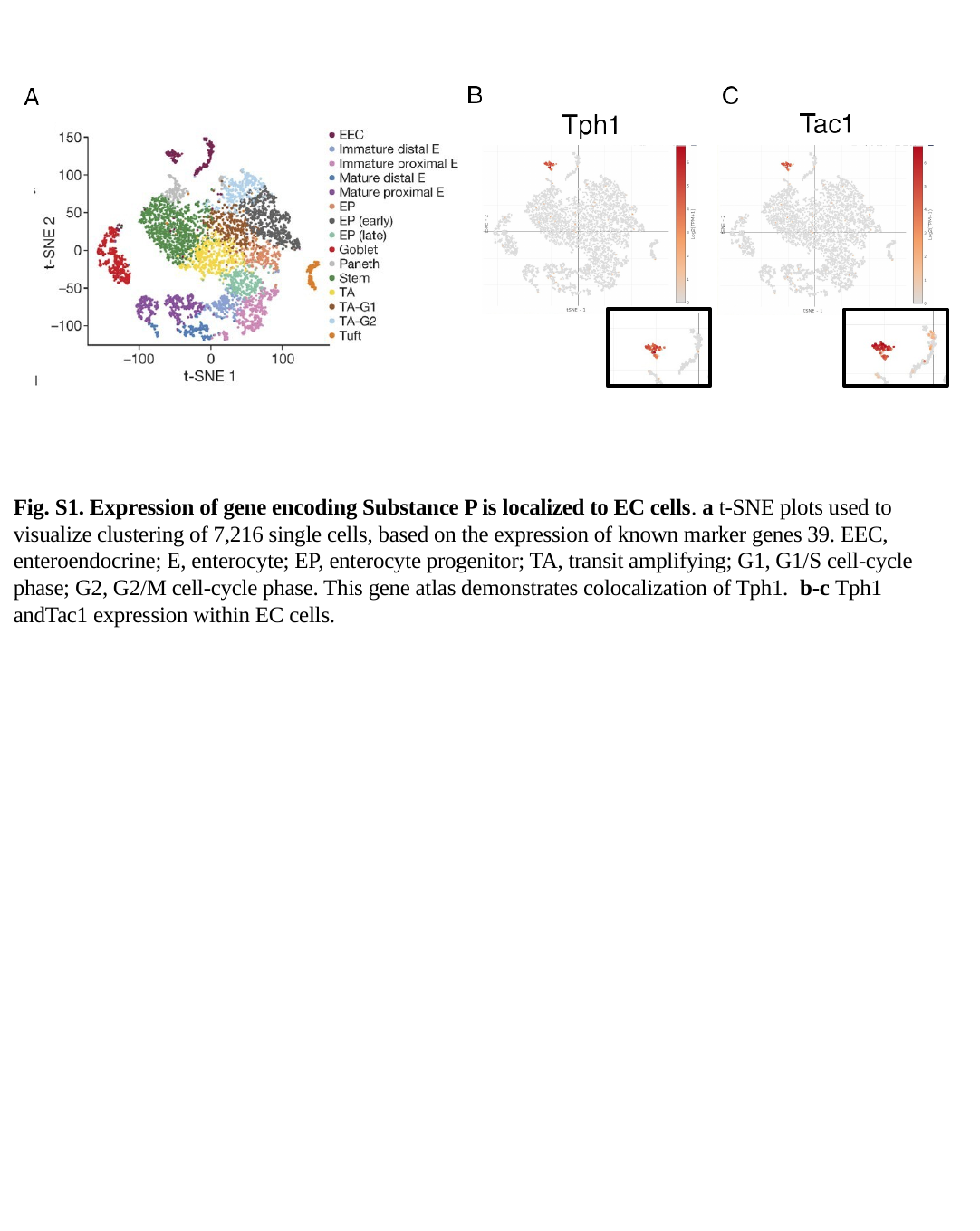

Fig. S1. Expression of gene encoding Substance P is localized to EC cells. a t-SNE plots used to visualize clustering of 7,216 single cells, based on the expression of known marker genes 39. EEC, enteroendocrine; E, enterocyte; EP, enterocyte progenitor; TA, transit amplifying; G1, G1/S cell-cycle phase; G2, G2/M cell-cycle phase. This gene atlas demonstrates colocalization of Tph1. b-c Tph1 andTac1 expression within EC cells.

### Slide 2
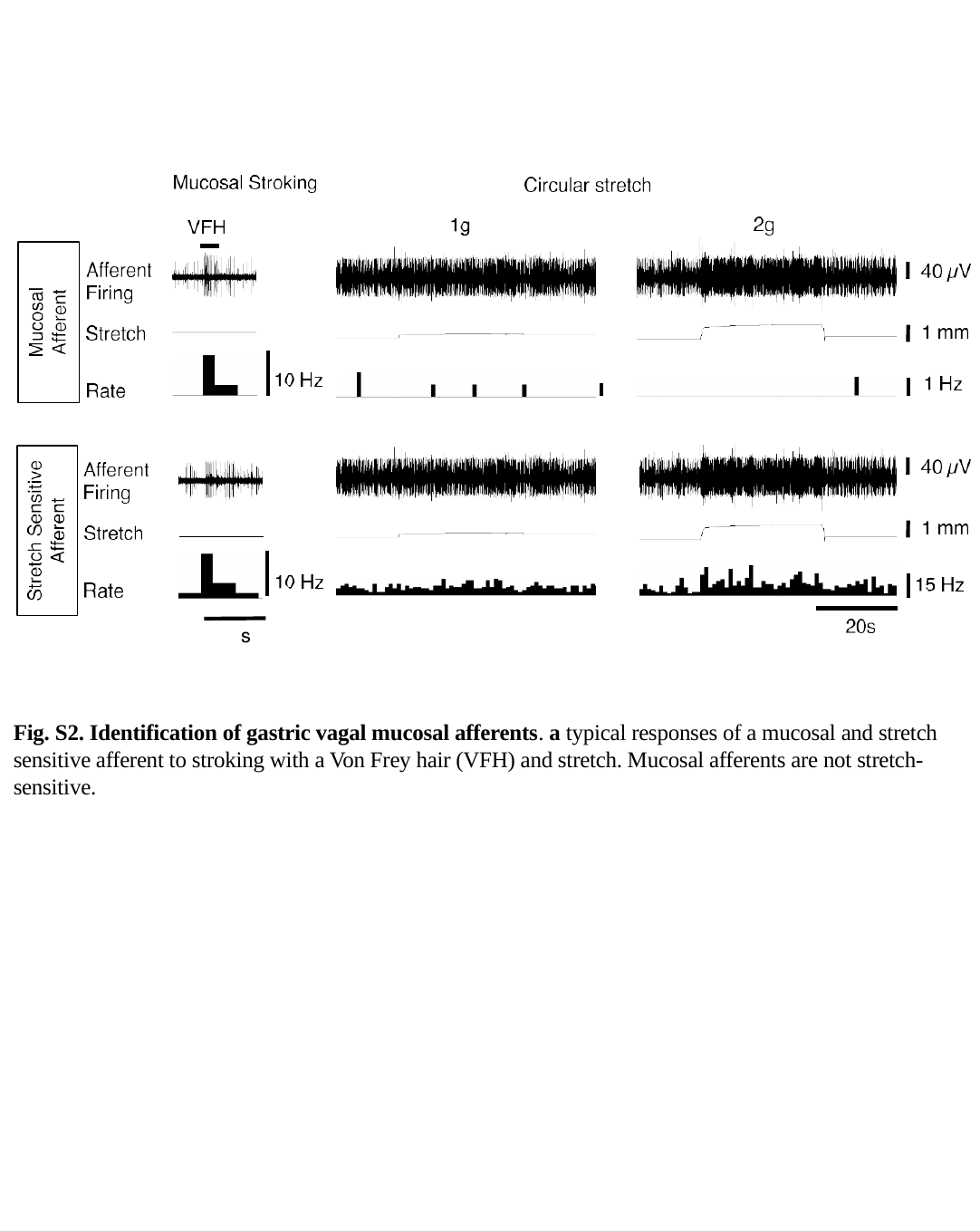

Fig. S2. Identification of gastric vagal mucosal afferents. a typical responses of a mucosal and stretch sensitive afferent to stroking with a Von Frey hair (VFH) and stretch. Mucosal afferents are not stretch-sensitive.

### Slide 3
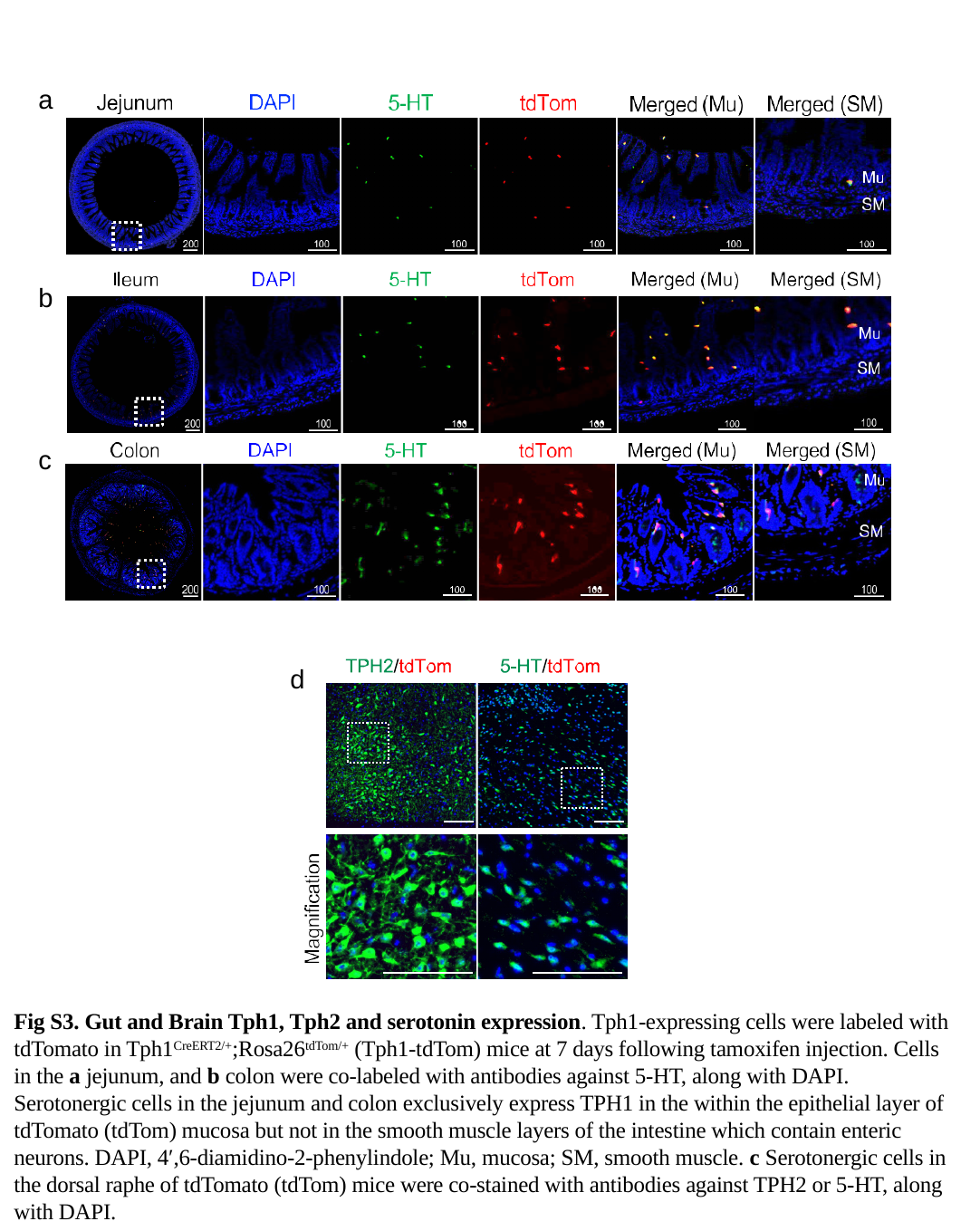

a
b
c
d
Fig S3. Gut and Brain Tph1, Tph2 and serotonin expression. Tph1-expressing cells were labeled with tdTomato in Tph1^CreERT2/+^;Rosa26^tdTom/+^ (Tph1-tdTom) mice at 7 days following tamoxifen injection. Cells in the a jejunum, and b colon were co-labeled with antibodies against 5-HT, along with DAPI. Serotonergic cells in the jejunum and colon exclusively express TPH1 in the within the epithelial layer of tdTomato (tdTom) mucosa but not in the smooth muscle layers of the intestine which contain enteric neurons. DAPI, 4′,6-diamidino-2-phenylindole; Mu, mucosa; SM, smooth muscle. c Serotonergic cells in the dorsal raphe of tdTomato (tdTom) mice were co-stained with antibodies against TPH2 or 5-HT, along with DAPI.

### Slide 4
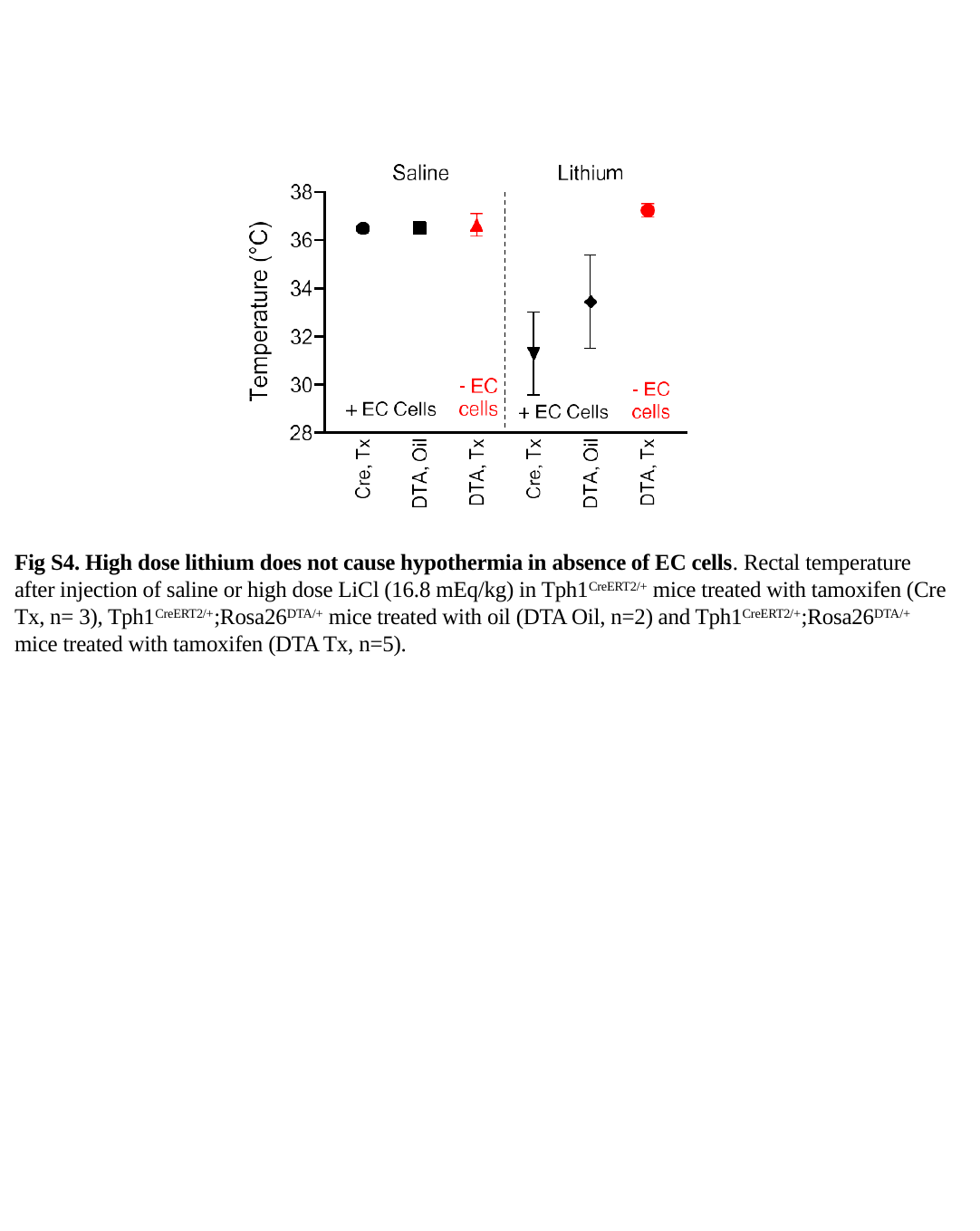

Fig S4. High dose lithium does not cause hypothermia in absence of EC cells. Rectal temperature after injection of saline or high dose LiCl (16.8 mEq/kg) in Tph1^CreERT2/+^ mice treated with tamoxifen (Cre Tx, n= 3), Tph1^CreERT2/+^;Rosa26^DTA/+^ mice treated with oil (DTA Oil, n=2) and Tph1^CreERT2/+^;Rosa26^DTA/+^ mice treated with tamoxifen (DTA Tx, n=5).

### Slide 5
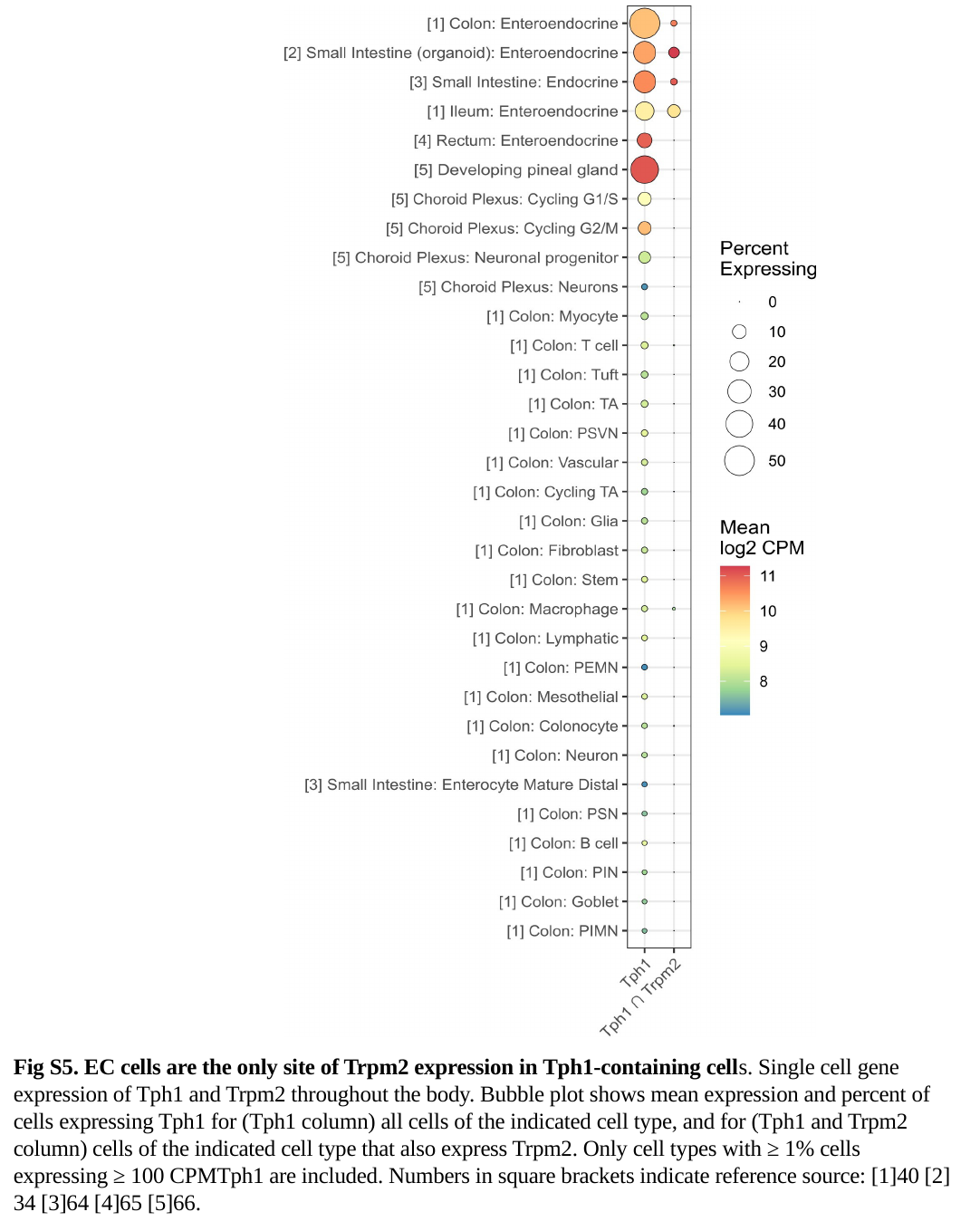

Fig S5. EC cells are the only site of Trpm2 expression in Tph1-containing cells. Single cell gene expression of Tph1 and Trpm2 throughout the body. Bubble plot shows mean expression and percent of cells expressing Tph1 for (Tph1 column) all cells of the indicated cell type, and for (Tph1 and Trpm2 column) cells of the indicated cell type that also express Trpm2. Only cell types with ≥ 1% cells expressing ≥ 100 CPMTph1 are included. Numbers in square brackets indicate reference source: [1]40 [2] 34 [3]64 [4]65 [5]66.

### Slide 6
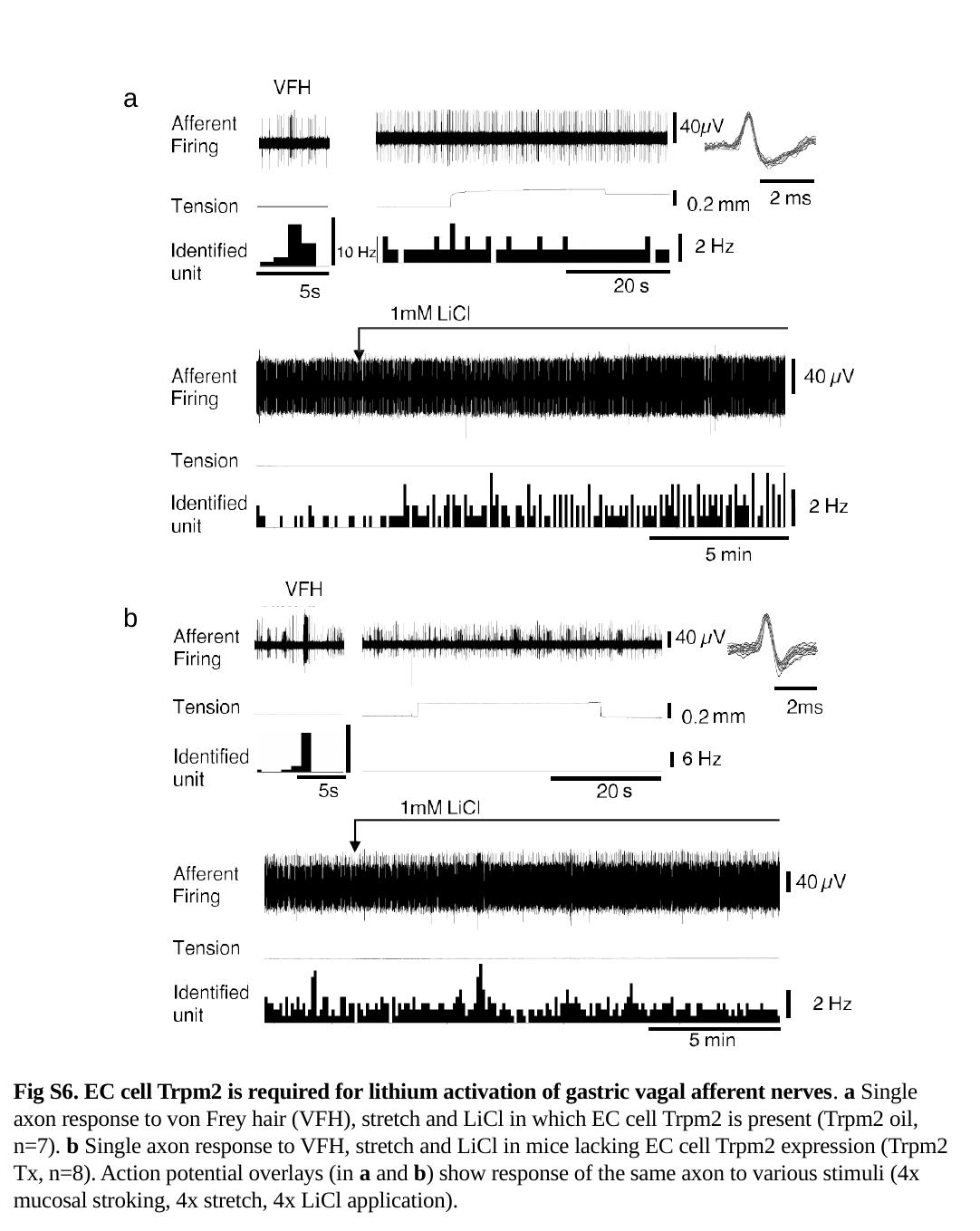

a
b
Fig S6. EC cell Trpm2 is required for lithium activation of gastric vagal afferent nerves. a Single axon response to von Frey hair (VFH), stretch and LiCl in which EC cell Trpm2 is present (Trpm2 oil, n=7). b Single axon response to VFH, stretch and LiCl in mice lacking EC cell Trpm2 expression (Trpm2 Tx, n=8). Action potential overlays (in a and b) show response of the same axon to various stimuli (4x mucosal stroking, 4x stretch, 4x LiCl application).

### Slide 7
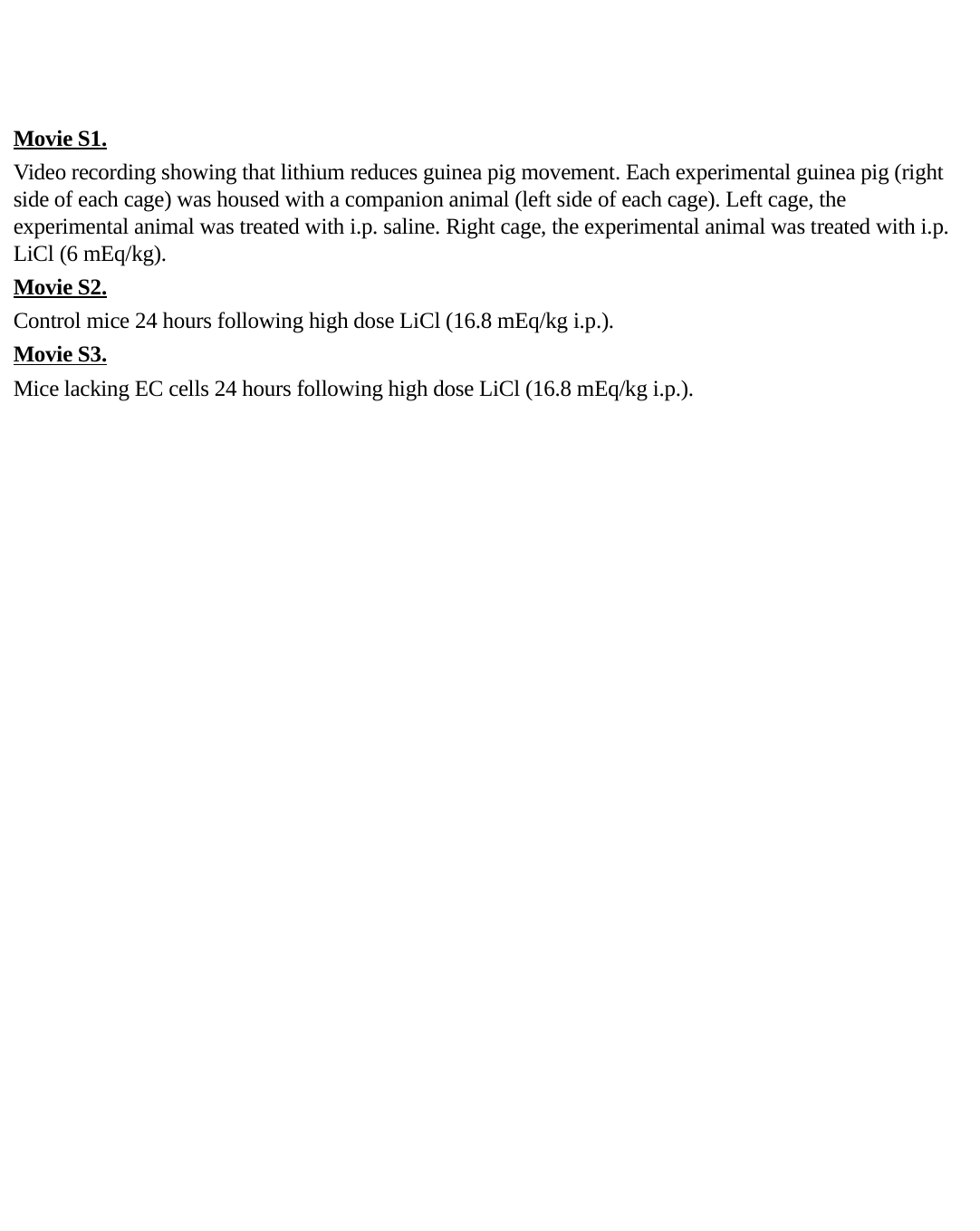

Movie S1.
Video recording showing that lithium reduces guinea pig movement. Each experimental guinea pig (right side of each cage) was housed with a companion animal (left side of each cage). Left cage, the experimental animal was treated with i.p. saline. Right cage, the experimental animal was treated with i.p. LiCl (6 mEq/kg).
Movie S2.
Control mice 24 hours following high dose LiCl (16.8 mEq/kg i.p.).
Movie S3.
Mice lacking EC cells 24 hours following high dose LiCl (16.8 mEq/kg i.p.).
